# A designed overlapping variant immunogen pool elicits broad sarbecovirus neutralization

**DOI:** 10.64898/2026.06.03.729821

**Authors:** Trinity Zang, Viren A. Baharani, Miranda Aldis, Marie Canis, Rachel Patejak, Edmund Osei Kuffour, Hans-Heinrich Hoffman, Harm van Bakel, Emilia M. Sordillo, Viviana Simon, Margaret R. MacDonald, Charles M. Rice, Michel C. Nussenzweig, Theodora Haziionannou, Paul D. Bieniasz

## Abstract

A central problem in achieving vaccine-based protection against viral infections is eliciting antibodies that are resilient to viral variation. Successive waves of SARS-CoV-2 infection during the COVID19 pandemic were driven by variants that acquired resistance to neutralizing antibodies elicited by prior SARS-CoV-2 variants. To the extent that serum neutralization breadth occurs in individuals with multiple exposures to SARS-CoV-2 antigens, we and others find that it is largely comprised of antibodies that target the variable receptor binding domain (RBD), rather than more conserved spike protein domains. By designing synthetic dimeric RBD immunogens we show that limiting divergence in heterodimeric components favors the generation of cross-reactive B cells and antibodies. We thus devised a vaccine approach based on a two-dose immunization with a pool of five overlapping heterodimeric synthetic RBD variants. Collectively, the RBD heterodimer pool was designed to cover 10% sequence variation and elicited greater antibody cross-reactivity and neutralization breadth than homodimers or heterodimers with highly divergent components. Using an unconventional ‘prospective’ challenge model in mice, we demonstrate the effectiveness of the RBD heterodimer pool in inducing antibody responses that attenuate infection by future SARS-CoV-2 variants, as well as protection in a challenge model based on a chimeric vesicular stomatitis virus bearing a spike protein from SARS-CoV-1.

**Significance statement:** Viral antigenic escape undermines both vaccination efforts and the development of herd immunity, resulting in an enormous viral disease burden. A central problem in eliciting vaccine-based protection against some viral infections is achieving antibody neutralization breadth. To elicit collections of antibodies that overcome the problem of limited antibody tolerance of viral variation, we designed a novel strategy based on an overlapping series of immunogens. This immunogen pool conferred at least partial protection against subsequently prevalent SARS-CoV-2 variants as well as a chimeric SARS-CoV-1 based model virus.

## Introduction

Viral antigenic escape occurs in large part through the acquisition of amino acid substitutions that attenuate the binding of neutralizing antibodies. The propensity of viruses to escape antibodies in this manner facilitates their persistence within individuals and in human populations. Viral antigenic escape undermines both vaccine development efforts and the development of herd immunity, resulting in an enormous viral disease burden(1). Viruses that are unable to escape antibody neutralization in this manner, such as measles virus, can theoretically be eradicated from humans through vaccination(2), while those that have great capacity for antigenic escape, such as human immunodeficiency virus type I and hepatitis C virus, cannot currently be vaccinated against(3). A central problem, therefore, in eliciting vaccine-based protection against some viral infections is achieving antibody neutralization breadth.

Successive waves of SARS-CoV-2 infection during the COVID19 pandemic were driven by novel viral variants that differed from the ancestral Wuhan-Hu-1 SARS-CoV-2 strain (SARS-CoV-2_Wu_)(4). New variants had constellations of substitutions that conferred resistance to neutralizing antibodies elicited by infection or vaccination with prior SARS-CoV-2 variants(5–10). Studies early in the pandemic revealed that the majority of SARS-CoV-2 neutralizing antibodies in the sera of previously infected or vaccinated individuals are directed against the receptor binding domain (RBD) and the N-terminal domain (NTD) of the spike protein(8, 11–16). These two portions of the spike protein became the most variable sequences in the SARS-CoV-2 genome(4, 17), as a consequence of selection pressure imposed by neutralizing antibodies(5–10), and despite (in the case of the RBD) constraints imposed by the need to interact with the ACE2 receptor(18).

A few individual SARS-CoV-2 neutralizing antibodies have been isolated that have greater breadth than is typical of RBD binding antibodies. These antibodies target more conserved portions of the spike protein, including conserved epitopes in the RBD, or other portions of spike such as those involved in membrane fusion(19–28). However, it is not clear whether the breadth that can sometimes be observed in human sarbecovirus neutralizing sera is based on antibodies that target non-RBD portions of the spike protein.

One promising vaccine strategy to elicit antibodies with greater breadth is the use of heteromultimeric or “mosaic” immunogens. In this approach, collections of related antigens, such as diverse sarbecovirus RBD domains, are displayed on the surface of nanoparticles(29–34). The theoretical rationale for this approach is that heteromultimeric antigens preferentially stimulate B cells whose B cell receptors (BCR) can be cross-linked by more than one related and physically linked antigen. Such cross-reactive BCRs would be predicted to give rise to cross-reactive, and ultimately broadly neutralizing, antibodies. Indeed, some examples of broadly neutralizing RBD-binding monoclonal antibodies have been cloned from animals experimentally immunized with mosaic nanoparticles(35–37). However, it is not clear what fraction of the antibodies elicited by mosaic immunogens are cross-reactive, and whether it is cross-reactive antibodies that are responsible for protection in vaccination experiments, which generally involve challenges with viruses genetically close to at least one component of the mosaic vaccine.

In our previous work, we analyzed serum antibodies in mice immunized with heterodimeric RBD immunogens, composed of two linked but diverse sarbecovirus RBDs, from SARS-CoV-1 and SARS-CoV-2_Wu_, or from SARS-CoV-2_Wu_ and SARS-CoV-2_BA.5_(38). While immunized animals readily developed antibodies that bound each individual component of an RBD-RBD heterodimer, most neutralizing antibodies bound just one, not both, RBD components of the heterodimer(38). Thus, the polyclonal antibodies elicited by RBD heterodimers include two largely separate sets of neutralizing antibodies that target each RBD component separately, not cross-reactive broadly neutralizing antibodies that bind both RBDs.

Notably, variation among sarbecovirus RBD sequences is concentrated in the neutralizing epitopes, presumably as a consequence of selection pressure being focused on these sequences. Extensive divergence in the neutralizing epitopes of an RBD heteromultimer decreases the likelihood that a naive BCR precursor of a potential neutralizing antibody could be cross-linked by RBD heteromultimers(38). Moreover, in our prior work we found that single amino acid substitutions are usually sufficient to confer resistance to most RBD binding neutralizing antibodies(5, 7, 39). Even ‘broadly’ neutralizing RBD specific antibodies (those that could neutralize SARS-CoV-2_Wu_, SARS-CoV-2_BA.1_ and SARS-CoV-2_BA.2_) could be escaped by 1 or 2 substitutions(10). This result suggests that there are stringent limits on the degree to which most antibodies (and by inference BCRs) can tolerate variation within an epitope. Thus, the initial B cell response to divergent heteromultimeric RBD immunogens may be biased toward conserved, but mostly non-neutralizing, epitopes.

Herein, we show that the neutralizing breadth encountered in human sera following repeated exposure to homologous and heterologous SARS-CoV-2 antigens is mostly directed at RBD epitopes, and natural SARS-CoV-2 variants escape these antibodies through loss of binding. To elicit collections of antibodies with greater breadth and to overcome the problem of limited antibody tolerance of epitope variation, while covering extensive sequence space, we designed an overlapping series of five heterodimeric RBD immunogens. Importantly, the level of epitope variation within each heterodimer was limited, to increase the probability of generating cross-reactive antibodies. This immunogen pool generated B cells and antibodies that displayed greater cross-reactivity than conventional heterodimers with highly divergent RBD components and conferred at least partial protection against subsequently prevalent SARS-CoV-2 variants as well as a chimeric VSV/SARS-CoV-1 construct.

## Results

### Neutralization breadth to SARS-CoV-2 variants conferred by RBD specific antibodies

Previously, we and others showed that the omicron BA.1 strain of SARS-CoV-2 represented a major antigenic shift and was largely resistant to neutralization by antibodies initially elicited by infection or vaccination by the ancestral SARS-CoV-2_Wu_ strain(40–42). However, sera from individuals who had received multiple exposures to ancestral antigen or breakthrough infection by SARS-CoV-2_BA.1_ displayed greater neutralizing breadth than sera from individuals who had been infected once by the ancestral SARS-CoV-2_Wu_ variant, or only received a 2-dose ancestral SARS-CoV-2_Wu_ mRNA vaccine(40, 43, 44). Previous studies of SARS-CoV-2_Wu_ vaccine recipient sera have suggested that RBD-directed antibodies dominate the neutralizing response against homologous SARS-CoV-2_Wu_ and heterologous SARS-CoV-2_BA.1_(12). Moreover, while individual RBD specific antibodies that neutralized SARS-CoV-2_Wu_ and SARS-CoV-2_BA.1_ could be isolated from individuals with multiple antigen exposures(10), the vast majority of these antibodies were escaped by substitutions(10) that appeared in subsequent variants with more extensive changes that emerged later in the pandemic (such as SARS-CoV-2_XBB_ which has substitutions at 21 out of 204 (10.3%) RBD amino acid positions compared to SARS-CoV-2_Wu_). Prior to this study, it was not clear whether the residual cross-reactive ability of polyclonal antibodies with subsequent variants to which individuals had not yet been exposed was conferred only by RBD antibodies, or whether antibodies targeting more conserved portions of spike(19–26) might also contribute. To determine which portion of SARS-CoV-2_Wu_ and SARS-CoV-2_BA.1_ spike elicited neutralizing antibodies in humans capable of cross-reacting with SARS-CoV-2_Wu_, SARS-CoV-2_BA.1_ and SARS-CoV-2_XBB_, we conducted plasma depletion studies. Individuals with multiple antigen exposures, including two groups that had been exposed to ancestral SARS-CoV2_Wu_-like antigens only: (i) those infected with ancestral SARS-CoV2_Wu_, then vaccinated with the SARS-CoV2_Wu_-based Pfizer–BioNTech or Moderna mRNA vaccines(ConVax, n=15) and (ii) those vaccinated with three doses of Pfizer–BioNTech or Moderna mRNA vaccines (Vax3, n=15). A third group had been vaccinated with three doses of Pfizer–BioNTech mRNA and then had exposure to a heterologous antigen via breakthrough SARS-CoV2_BA.1_ infection (Breakthrough, or BT, n=13).

Neutralizing titers against SARS-CoV-2_Wu_, SARS-CoV-2_BA.1_ and SARS-CoV-2_XBB_ pseudotypes were determined for sera from all individuals (Fig.1A). Sera from all individuals neutralized SARS-CoV-2_Wu_, with mean NT50 of 30,524 (ConVax), 7,674 (Vax3), and 15,660 (Breakthrough). SARS-CoV-2_BA.1_ was also neutralized (mean NT50 = 2,680 ConVax, 1,245 Vax3, and 4,530 Breakthrough), representing levels that were a mean of 14%, 22% and 37% of the SARS-CoV-2_Wu_ neutralizing activity. These sera had only low neutralizing activity against SARS-CoV-2_XBB_, (mean NT_50_ = 184 ConVax, 88 Vax3, and 259 Breakthrough) (Fig.1A).

**Fig. 1.**
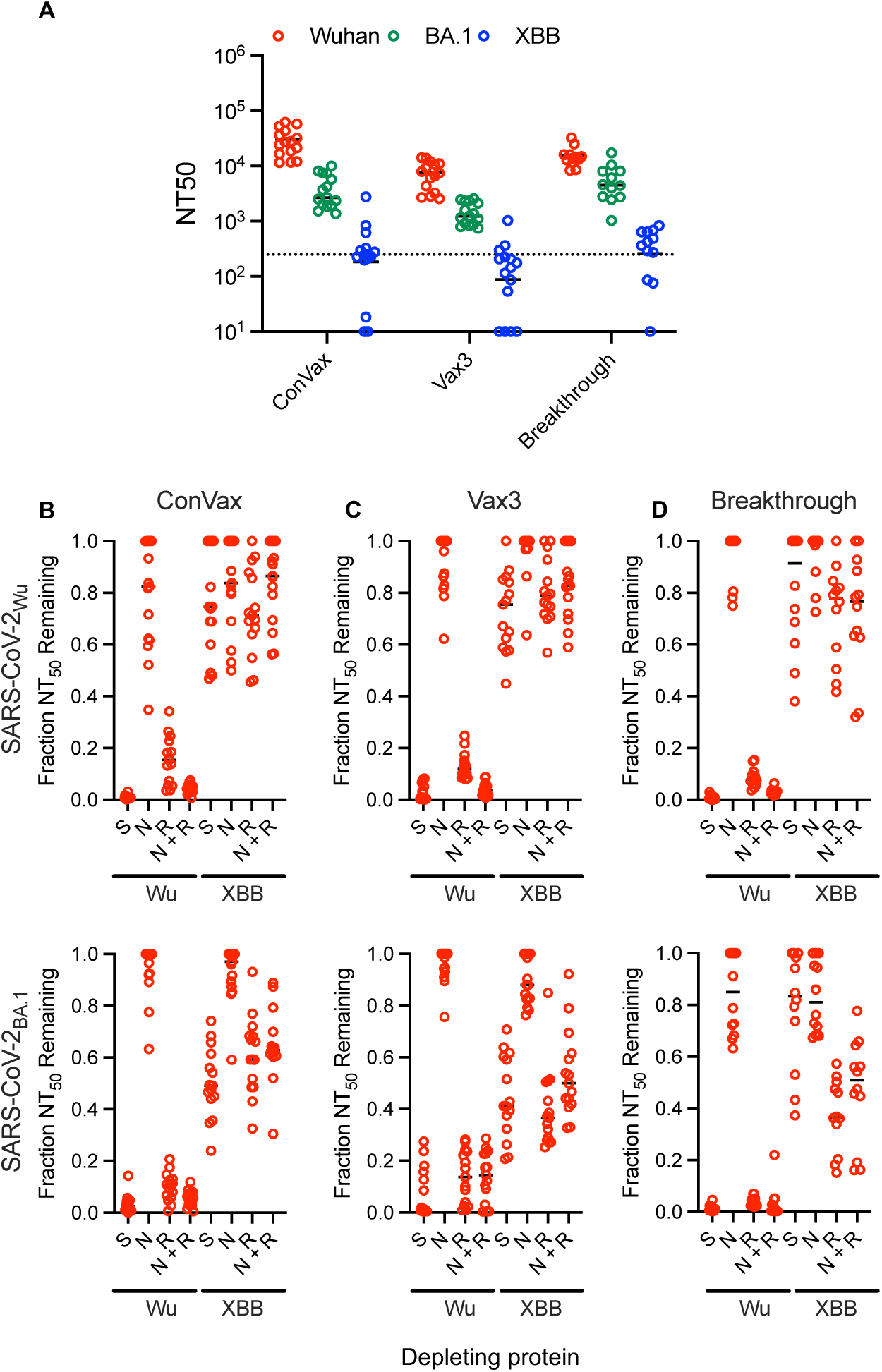
Neutralizing breadth following SARS-CoV-2 infection and or vaccination is conferred by RBD-specific antibodies. (A) Neutralizing titers (NT_50_) against SARS-CoV-2 pseudotype variants in ConVax, Vax3, and Breakthrough patient plasma. Each symbol represents 1 participant, lines = group mean. ConVax n=15, Vax3 n=15, Breakthrough n=13. Dotted line indicates the lowest sera dilution tested (1:250). (B-D) Fraction of neutralizing titers against SARS-CoV-2_Wu_ and SARS-CoV-2_BA.1_ pseudotypes remaining in ConVax(B), Vax3(C), and Breakthrough(D) patient plasma following depletion by the indicated Spike(S), NTD(N), RBD(R), or mixed NTD and RBD(N+R) protein. Each symbol represents one participant, lines=group mean. ConVax n=15, Vax3 n=15, Breakthrough n=13.

To determine which domains of spike elicited the plasma antibodies that neutralized SARS-CoV-2_Wu_, and SARS-CoV-2_BA.1_ in each group, we incubated plasma with 6xHis-tagged recombinant proteins comprising full length Spike_Wu_ (S), or the isolated NTD_Wu_ (N), or RBD_Wu_ (R) domains, alone or in combination (N + R). Protein-antibody complexes were depleted with His-tagged Dynabeads and remaining neutralizing activity against SARS-CoV-2_Wu_ and SARS-CoV-2_BA.1_ was measured (Fig.1B-D, Fig.S1A-C). The full length Spike_Wu_ depleted nearly all the neutralizing activity against SARS-CoV-2_Wu_ (mean = 99%, 97%, and 99% for the ConVax, 3xVax and BT groups respectively) (Fig.1B-D, Fig.S1A-C). Antibodies targeting the NTD and RBD accounted for nearly all of the neutralizing activity, as the mixture of NTD and RBD protein depleted 95% (ConVax), 96% (3xVax) and 97% (BT) of the neutralizing activity similarly to full length spike (Fig.1B-D, Fig.S1A-C). In fact, most of the neutralizing activity was depleted by RBD_Wu_ alone (84% (ConVax), 86% (3xVax) and 91% (BT) and a comparatively minor fraction was depleted by the NTD_Wu_, (21% (ConVax), 8% (3xVax) and 6% (BT) (Fig.1B-D, Fig.S1A-C). For heterologous neutralization of SARS-CoV-2_BA.1_, RBD_Wu_ alone similarly depleted 90% (ConVax), 86% (3xVax) and 96% (BT) of the neutralizing activity with a small fraction depleted by the NTD_Wu_, (6% (ConVax), 4% (3xVax) and 15% (BT) (Fig.1B-D, Fig.S1A-C). Overall, the individuals who had received multiple antigen exposures had neutralizing antibodies that cross-reacted with SARS-CoV-2_BA.1_ and the majority of these cross-reactive neutralizing antibodies were directed against the RBD, as evidenced by the ability of RBD_Wu_ to deplete the activity against SARS-CoV-2_BA.1_.

We repeated neutralization-depletion experiments using recombinant proteins from the more genetically distant variant SARS-CoV-2_XBB_ (Fig.1B-D, Fig.S1D-F). Neutralizing antibodies against ancestral SARS-CoV-2_Wu_ were largely resistant to depletion by SARS-CoV-2_XBB_-derived proteins. However, some of the cross-reactive neutralizing activity against SARS-CoV-2_BA.1_ was depleted by SARS-CoV-2_XBB_ spike (mean = 50% (ConVax), 55% (3xVax) and 21% (BT)). The RBD_XBB_ (but not the NTD_XBB_) was as effective as the full-length spike in depleting these neutralizing antibodies (mean = 39% (ConVax), 60% (3xVax) and 62% (BT)) (Fig.1B-D, Fig. S1D-F). These results suggest that a small fraction of the neutralizing antibodies in these plasma cross-react with SARS-CoV-2_Wu_, SARS-CoV-2_BA.1_, and SARS-CoV-2_XBB_ and the vast majority of these cross-reactive neutralizing antibodies are directed against the RBD, not against the NTD or against other, more conserved portions of the spike protein, even after exposure to a heterologous antigen. This dominance of the RBD in eliciting cross reactive neutralizing antibodies after multiple or heterologous antigen exposures is consistent with recent findings of Torotorici et. al(45), and suggest that despite its intrinsic variability, the RBD should be the key component of vaccines designed to elicit broad neutralization against SARS-CoV-2 variants.

### Cross-reactive B cells and neutralizing antibodies induced by RBD heterodimers

In our previous work, immunization of mice with a heterodimeric RBD immunogen comprising of two divergent sarbecovirus or SARS-CoV-2 variant RBDs elicited polyclonal antibodies that contained two largely separate sets of single RBD variant-specific neutralizing antibodies, rather than cross-reactive broadly neutralizing antibodies that bind both RBDs(38). Reasoning that the extent of divergence between heterodimer components might impact their propensity to elicit cross-reactive versus single RBD-specific antibodies, we attempted to elicit cross-reactive neutralizing antibodies using more closely related heterodimer RBD pairs. We designed RBD homodimers and heterodimers encoding ancestral SARS-CoV-2_Wu_ and a synthetic RBD variant that we termed “Beta+”. Specifically, the Beta+ variant contains 5 substitutions relative to SARS-CoV-2_Wu_; three substitutions are from the natural SARS-CoV-2_Beta_ variant which contained the first bona fide antibody escape mutations in class I/II/III epitopes to emerge during the COVID19 pandemic (K417N, E484K, N501Y). The RBD_Beta+_ includes additional mutations located in the class III and class IV epitopes (R346T and S371F), spatially separated from the substitutions in RBD_Beta_. Thus RBD_Beta+_ differs from RBD_Wu_ at five spatially separated amino acid positions.

Groups of C57BL/6 mice were immunized at week 0 and 3 with (i) individual RBD homodimers (RBD_Wu_-RBD_Wu_, or RBD_Beta+_-RBD_Beta+_) (ii) a mixture of two homodimers (RBD_Wu_-RBD_Wu_ + RBD_Beta+_-RBD_Beta+_) or (iii) a heterodimer (RBD_Wu_-RBD_Beta+_). The total amount (mass) of RBD protein was kept constant for each immunization strategy. At 12 weeks after the first injection, mice were terminally bled and sera was assayed for neutralizing activity against a SARS-CoV-2_Wu_ pseudotype and a synthetic SARS-CoV-2_Beta+_ pseudotype whose RBD had the same 5 substitutions present in the RBD_Beta+_ immunogen (Fig. 2A,B). The RBD_Wu_-RBD_Wu_ homodimeric immunogen elicited neutralizing titers that were 14x more potent against SARS-CoV-2_Wu_ (mean NT_50_= 28,160) than against SARS-CoV-2_Beta+_ (mean NT_50_= 1,958). Surprisingly though, in the reciprocal case, the RBD_Beta+_-RBD_Beta+_ homodimer elicited neutralizing titers that were similar against SARS-CoV-2_Wu_ (mean NT_50_= 30,641) and SARS-CoV-2_Beta+_ (mean NT_50_= 14,544) (Fig. 2A,B). The mixed RBD_Wu_-RBD_Wu_ + RBD_Beta+_-RBD_Beta+_ homodimers elicited similar titers against SARS-CoV-2_Wu_ and SARS-CoV-2_Beta+_ (mean NT_50_= 16,309 vs 9,822 respectively) as did the RBD_Wu_-RBD_Beta+_ heterodimer (mean NT_50_= 30,234 vs 13,618 respectively) (Fig. 2A,B). The homodimers and heterodimers elicited modest titers of neutralizing antibodies against the more divergent SARS-CoV-2 strain (SARS-CoV-2_XBB_) (Fig. 2C). In this regard, the pooled homodimers RBD_Wu_-RBD_Wu_ + RBD_Beta+_-RBD_Beta+_ appeared similar to the RBD_Wu_-RBD_Beta+_ heterodimer (mean NT_50_= 322 vs 385 respectively) (Fig. 2C).

**Fig. 2.**
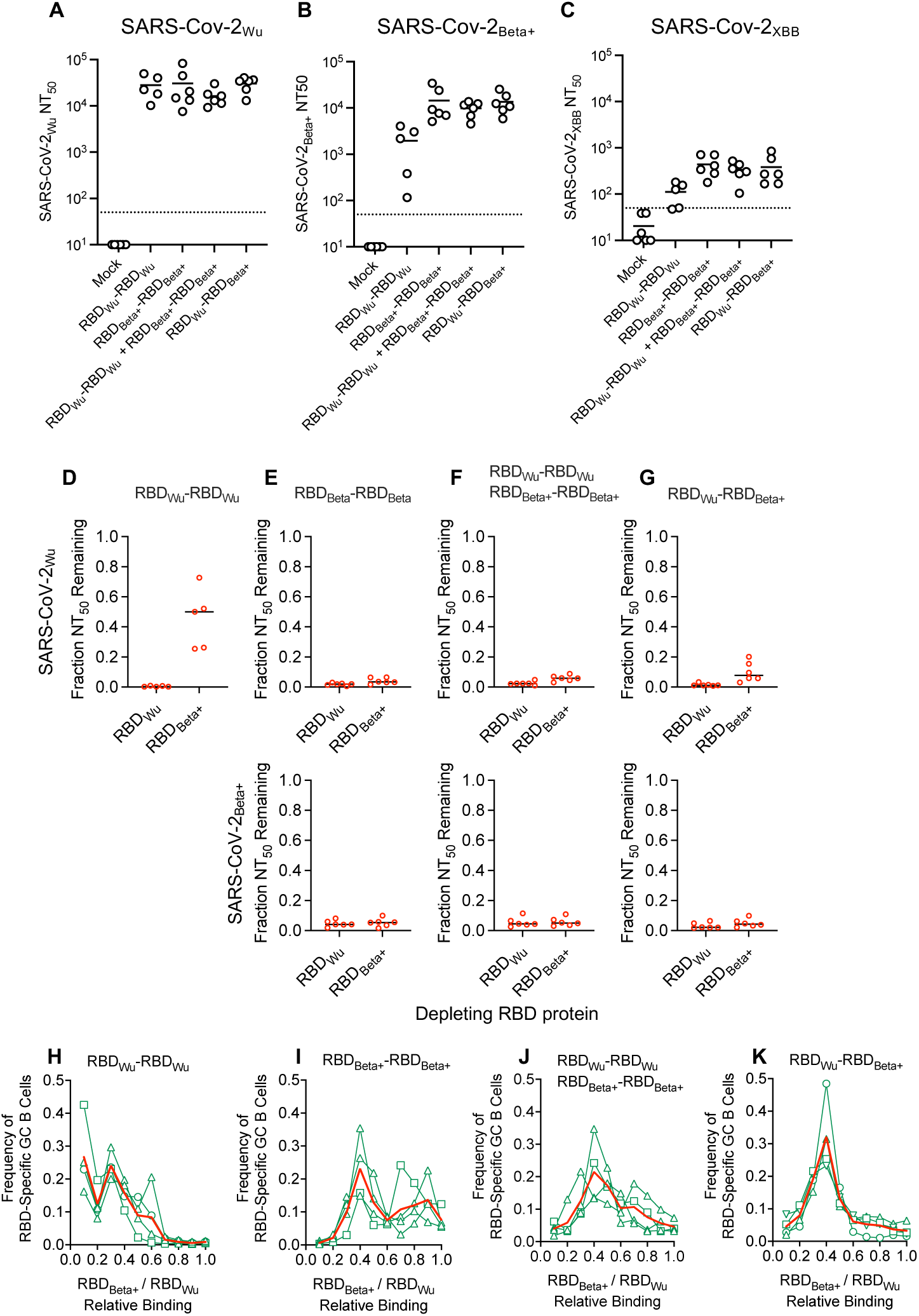
Cross-reactive neutralizing antibodies and B cells induced in homodimer and heterodimer immunized mice. (A-C) Neutralizing titers (NT_50_) against SARS-CoV-2 pseudotypes and variants thereof in mouse sera, 12 weeks post immunization with two doses of the indicated RBD dimer immunogens. Each symbol represents 1 mouse, lines = group mean, n = 4-6 mice per group. Dotted line indicates the lowest sera dilution tested (1:50). (D-G) Fraction of neutralizing titer against SARS-CoV-2_Wu_, or SARS-CoV-2_Beta+_ pseudotypes remaining in12 week post-immunization mouse sera after depletion with the indicated RBD protein. Graph title indicates immunogen group. Each symbol represents 1 mouse, lines represent group mean, n=4-6 mice per group. (H-K) Proportion of RBD-specific lymph node germinal center B cells (Y axis) that bind RBD_Wu_ and RBD_Beta+_ monomer baits at the indicated relative fluorescence intensity ratios (fluorescence intentity of RBD_Beta+_ binding divided by the sum of the fluorescence intensities of RBD_Wu_ and RBD_Beta+_ binding) plotted on the X-axis. Graph title indicates immunogen group. Each green line represents one mouse, red lines indicate the group mean (n=8).

We next used the RBD-6xHis depletion assay to test for cross-reactive neutralizing antibodies (Fig.2D-G, Fig.2SA-D). In sera from mice immunized with the RBD_Wu_-RBD_Wu_ homodimer, the neutralizing activity against SARS-CoV-2_Wu_ was completely depleted by RBD_Wu_, but only ∼50% of the neutralizing activity was depleted by RBD_Beta+_ (Fig. 2D). Neutralizing activity against SARS-CoV-2_Beta+_ elicited by RBD_Wu_-RBD_Wu_ was not sufficiently robust for conclusive results in the depletion assay (Fig. S2). However, neutralizing activity against SARS-CoV-2_Beta+_ elicited by RBD_Wu_-RBD_Wu_, or RBD_Beta+_-RBD_Beta+_ homodimers, or the RBD_Wu_-RBD_Beta+_ heterodimer could be depleted by RBD_Wu_ or RBD_Beta+_ (>90% depletion in each case) (Fig. 2D-G, Fig. S2). These results suggest that the RBD_Wu_-RBD_Wu_ homodimer elicits a mixture of RBD_Wu_-specific, and RBD_Wu_/RBD_Beta+_ cross-reactive antibodies while the RBD_Beta+_-RBD_Beta+_ homodimer and RBD_Wu_-RBD_Beta+_ heterodimer primarily elicit neutralizing antibodies that cross react with both RBD_Wu_ and RBD_Beta+_.

To further examine the B cells and antibodies elicited by the RBD_Wu_/RBD_Beta+_ homodimers and heterodimers, we injected mouse footpads with the homodimer and heterodimer immunogens and analyzed germinal center (GC) B-cells from the local draining popliteal lymph node for binding to RBD monomers. The relative fluorescent intensities of individual B-cells resulting from staining with a mixture of monomeric RBD_Wu_ and RBD_Beta+_ proteins labelled with different fluorophores was taken as a measure of the relative abilities of their B cell receptors to bind RBD_Wu_ and RBD_Beta+_ respectively (Fig. 2H-K, Fig. S3). In mice immunized with the RBD_Wu_-RBD_Wu_ homodimer, a sub-population of the RBD-specific GC B cells bound specifically to RBD_Wu_, while another portion of the B cells were cross-reactive, and bound to both RBD_Wu_ and RBD_Beta+_ (Fig. 2H, Fig. S3A). Notably, there was clear heterogeneity in the relative propensity of these cross-reactive B cells to bind RBD_Wu_ and RBD_Beta+_. Similarly, in mice immunized with the RBD_Beta+_-RBD_Beta+_ homodimer, cross-reactive binding to RBD_Wu_ and RBD_Beta+_ monomers was evident, with sub population-heterogeneity in the magnitude of binding to RBD_Wu_ and RBD_Beta+_ monomers, and smaller populations binding to RBD_Beta+_ alone (Fig. 2I, Fig. S3B). Likewise, in mice immunized with the mixture of RBD_Wu_-RBD_Wu_ and RBD_Beta+_-RBD_Beta+_ homodimers, heterogenous cross-reactive binding to RBD_Wu_ and RBD_Beta+_ monomers was evident with smaller populations binding to RBD_Wu_ or RBD_Beta+_ alone (Fig. 2J, Fig. S3). However, immunization with the RBD_Wu_-RBD_Beta+_ heterodimer almost exclusively elicited cross-reactive B cells that bound both RBD_Wu_- and RBD_Beta+_ monomers (Fig. 2K, Fig. S3D). Unlike the heterogenous cross-reactive binding B cells elicited by the RBD_Beta+_-RBD_Beta+_ and RBD_Wu_-RBD_Wu_/RBD_Beta+_-RBD_Beta+_ homodimers, the RBD_Wu_-RBD_Beta+_ heterodimer elicited comparatively homogenously cross-reactive B cells, with greater degree of concordance in their ability to bind to the RBD_Wu_ and RBD_Beta+_ monomers (Fig. 2K, Fig. S3D). Note that for the RBD_Wu_-RBD_Wu_/RBD_Beta+_-RBD_Beta+_ homodimer mix and the RBD_Wu_-RBD_Beta+_ heterodimer, the sequence composition of the immunizing proteins is effectively identical (50% RBD_Wu_ and 50% RBD_Beta+_) but with RBDs alternatively configured either as homodimers or heterodimers, with discernable effects on B cell cross reactivity. Together, these results suggest that an RBD heterodimer comprised of two closely related RBDs, such as RBD_Wu_-RBD_Beta+_, can elicit germinal center B cells that are mostly cross-reactive and bind more similarly to each dimer component than those elicited by individual homodimers or mixed homodimers.

### A designed synthetic variant pool of overlapping heterodimers elicits cross-reactive neutralizing antibodies

RBD heterodimers whose component RBDs are highly divergent failed to yield cross-reactive neutralizing antibodies(38). However, the aforementioned results indicate that it is possible to manipulate the B cell response to favor cross-reactive antibodies by using RBD heterodimers with more modest divergence. We therefore designed an overlapping pool of synthetic variant RBD heterodimers that would cumulatively encompass a high degree of diversity, including most of the naturally occurring variation that had been documented in SARS-CoV-2 at that time (Fig. 3A-D). However, the individual RBD-RBD heterodimer components of the synthetic pool were designed to exhibit modest intra-dimer variation. At the time of immunogen design (August 2023) the dominant circulating SARS-CoV-2 variant was SARS-CoV-2_XBB_, that contained substitutions at 21 out of 204 (10.3%) RBD amino acid positions compared to the ancestral SARS-CoV-2_Wu_ variant (Fig. 3B). Those substitutions concentrated in neutralizing epitopes. The RBD heterodimer pool was composed of a series of five overlapping heterodimers, where the C-terminal RBD of each dimer in the set was identical to the N-terminal RBD of the next heterodimer in the overlapping series, beginning with the aforementioned RBD_Wu_-RBD_Beta+_ heterodimer. In this first component of the pool, the two RBDs differed at 5 amino acid positions and in each subsequent heterodimer four or five additional substitutions were added in a stepwise manner to the C-terminal RBD. The substitutions at each step were selected to be spatially separated and distributed among the various neutralizing epitope classes(46) (Fig. 3A-D). The resulting ovelapping dimer pool termed “Wuhan-to-XBB x 5” (WX5) was comprised of five heterodimers: (i) RBD_Wu_-RBD_Beta+_, (ii) RBD_Beta+_-RBD_Beta2+_ (iii) RBD_Beta2+_-RBD_Beta3+_ (iv) RBD_Beta3+_-RBD_Beta4+_ (v) RBD_Beta4+_-RBD_XBB_) (Fig. 3A,D). We also constructed an RBD_Wu_-RBD_XBB_ heterodimer for comparative purposes, that contained the same 21 amino acid substitutions as the RBD_WX5_ set, but with all substitutions relative to RBD_Wu_ contained in the C-terminal RBD_XBB_ component of a single heterodimer.

**Fig. 3.**
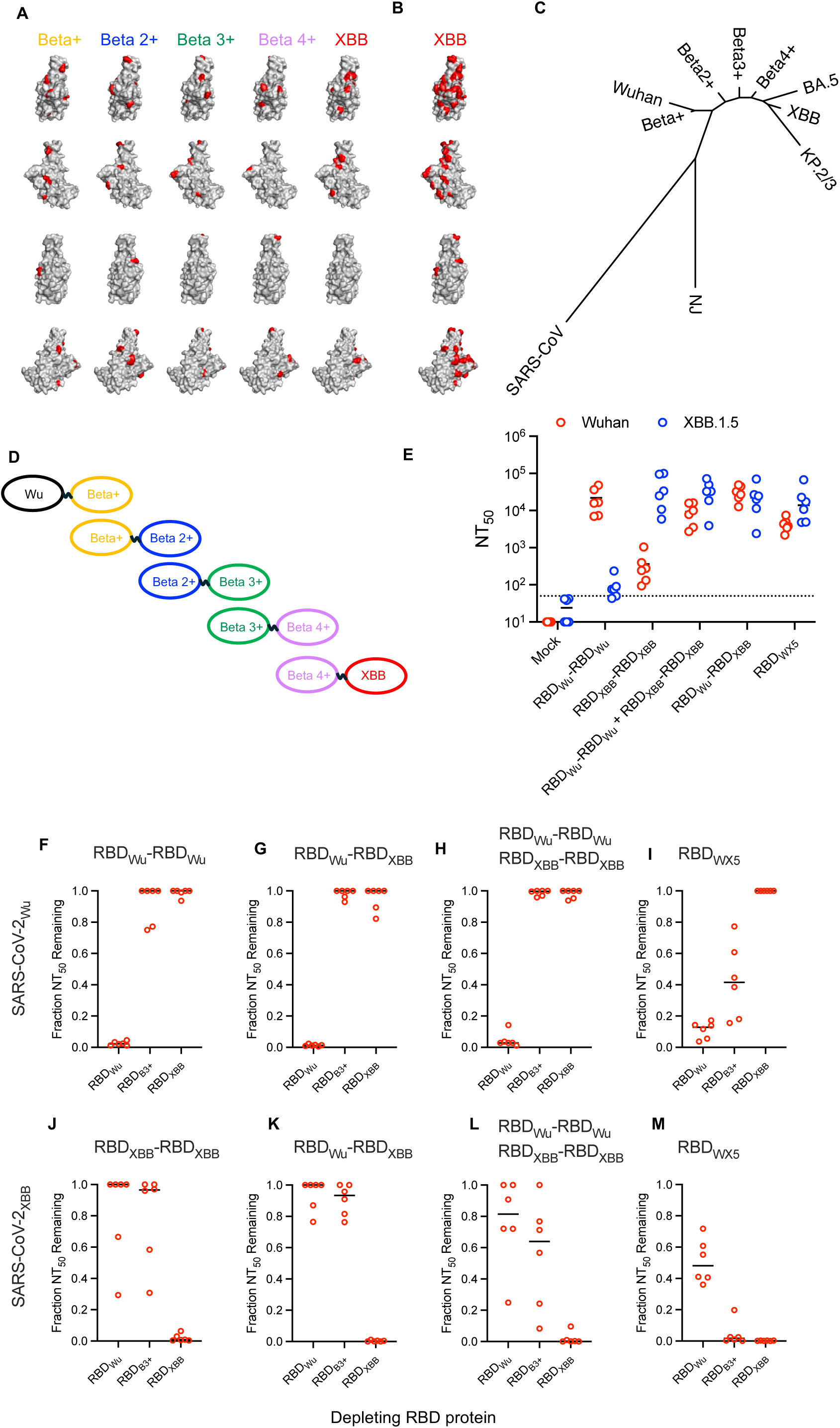
A pool of closely related overlapping synthetic RBD heterodimers elicits broadly neutralizing antibodies. (A) Components of the overlapping RBD_WX5_ pool. SARS-CoV-2 RBD structure with the amino acids colored red that are changed in each progressive iteration from the previous RBD in the series (Wuhan to Beta+, Beta+ to Beta2+, Beta2+ to Beta3+, Beta3+ to Beta4+, Beta4+ to XBB). Each row represents a view of the RBD rotated by 90° about the vertical axis from the view above. (B) Accumulated changes in the XBB strain relative to Wuhan, which represent the extreme ends of the RBD_WX5_ series. (C) Neighbor joining tree constructed using the amino acid sequences of the RBD from the Wuhan, XBB and synthetic RBD components of the RBD_WX5_ pool as well as RBD sequences from naturally occurring SARS-CoV2 variants and SARS-CoV. (D) Schematic representation of the 5 heterodimer components of the RBD_WX5_ immunogen. (E) Neutralizing titers (NT_50_) against SARS-CoV-2_Wu_ and SARS-CoV-2_XBB_ pseudotypes in mouse sera at 12 weeks post immunization with two doses of the indicated RBD dimer immunogens. Each symbol represents 1 mouse, lines represent group mean, n=6 mice per group. Dotted line indicates the lowest serum dilution tested (1:50). (F-I) Fraction of neutralizing titer (NT_50_) against SARS-CoV-2_Wu_ pseudotypes remaining in 12 week post-immunization mouse sera from mice immunized with RBD_Wu_-RBD_Wu_ homodimer (F) RBD_Wu_-RBD_XBB_ heterodimer (G) a mixture of the RBD_Wu_-RBD_Wu_ and RBD_XBB_-RBD_XBB_ homodimers (H), and the RBD_WX5_ heterodimer pool (I) after depletion with the indicated RBD proteins. (J-M). Fraction of neutralizing titer (NT_50_) against SARS-CoV-2_XBB_ pseudotypes remaining in 12 week post-immunization mouse sera from mice immunized with RBD_XBB_-RBD_XBB_ homodimer (J), RBD_Wu_-RBD_XBB_ heterodimer (K), a mixture of the RBD_Wu_-RBD_Wu_ and RBD_XBB_-RBD_XBB_ homodimers (L), and the RBD_WX5_ pool (M) after depletion with the indicated RBD proteins For F-M, graph title indicates immunogen group, each symbol represents 1 mouse, lines indicate group mean, n=6 mice per group.

Five groups of K18-hACE2 mice were immunized at week 0 and 3, with individual (i) RBD_Wu_-RBD_Wu_, or (ii) RBD_XBB_-RBD_XBB_ homodimers, (iii) a 1:1 mixture of RBD_Wu_-RBD_Wu_ + RBD_XBB_-RBD_XBB_, homodimers, (iv) the RBD_Wu_-RBD_XBB_ heterodimer or (v) the RBD_WX5_ heterodimer overlapping pool. The total amount of immunogen was fixed at 10 μg and mice were bled at week 0, 3, 7 and 12 to measure serum neutralizing activity against SARS-CoV-2_Wu_ and SARS-CoV-2_XBB_ (Fig. 3E, Fig. S4A-E). The RBD_Wu_-RBD_Wu_ homodimer generated high titer antibodies against SARS-CoV-2_Wu_ (mean NT50 = 21,889) and very low titer against SARS-CoV-2_XBB_ (mean NT50 = 95), and conversely the RBD_XBB_-RBD_XBB_ homodimer generated high titer antibodies against SARS-CoV-2_XBB_ (mean NT50 = 45,018) and only modest titer against SARS-CoV-2_Wu_ (mean NT50 = 364) (Fig. 3E, Fig. S4A,B). The mixed RBD_Wu_-RBD_Wu_ + RBD_XBB_-RBD_XBB_ homodimers, the RBD_Wu_-RBD_XBB_ heterodimer and the RBD_WX5_ heterodimer pool all generated high titer antibodies that neutralized SARS-CoV-2_Wu_ and SARS-CoV-2_XBB_ (mean NT50 = 4,353 to 34,926) (Fig. 3E, Fig. S4C-E).

To test for the presence of cross-reactive neutralizing antibodies we depleted 12-week sera with RBD_Wu_ or RBD_XBB_ monomers (that differ at 21 positions from each other) or with RBD_Beta3+_, one component of the heterodimer set that differs at 15 amino acid positions from RBD_Wu_ and at 10 positions from RBD_XBB_. We then measured the remaining neutralizing activity against SARS-CoV-2_Wu_ and SARS-CoV-2_XBB_ (Fig. 3F-M, Fig. S5A-E). As expected from our earlier studies with homodimers and genetically distant RBD heterodimers, RBD_Wu_-RBD_Wu,_ RBD_XBB_-RBD_XBB_, and RBD_Wu_-RBD_XBB_ generated separate sets of non cross-reactive SARS-CoV-2_Wu_ and SARS-CoV-2_XBB_ neutralizing antibodies, that in most cases were only depleted by the homologous RBD (Fig. 3F,G,J,K, Fig. S5A-C). The pooled homodimers, RBD_Wu_-RBD_Wu_ + RBD_XBB_-RBD_XBB_, generated neutralizing antibodies against SARS-CoV-2_Wu_ that were not cross-reactive with

RBD_Beta3+_ or RBD_XBB_ (Fig. 3H, Fig. S5D), and neutralizing antibodies against SARS-CoV-2_XBB_ that were partly and inconsistently depleted by RBD_Wu_ (mean depletion = 23%) and RBD_Beta3+_ (mean depletion = 44%) in some mice (Fig. 3L, Fig. S5D). Conversely the RBD_WX5_ pool elicited SARS-CoV-2_Wu_ neutralizing antibodies that were partly cross-reactive with, and depleted by, RBD_Beta3+_ (mean depletion = 58%) (Fig. 3I, Fig. S5E), while the neutralizing antibodies against SARS-CoV-2_XBB_ were cross-reactive and could be nearly completely depleted by RBD_Beta3+_ (mean depletion = 96%) and partly depleted by RBD_Wu_. (mean depletion = 49%) (Fig. 3M, Fig. S5E). These data indicate that the SARS-CoV-2_Wu_ and SARS-CoV-2_XBB_ neutralizing antibodies elicited by the RBD_WX5_ heterodimer pool exhibited a higher degree of cross-reactivity than those elicited by the RBD_Wu_-RBD_XBB_ heterodimer and the pooled RBD_Wu_-RBD_Wu_ + RBD_XBB_-RBD_XBB_ homodimers.

We injected mouse footpads with the RBD_Wu_-RBD_Wu_ + RBD_XBB_-RBD_XBB_ homodimer mixture, the RBD_Wu_-RBD_XBB_ heterodimer or the RBD_WX5_ overlapping pool, and analyzed GC B-cells from the local draining lymph node. The RBD_WX5_ heterodimer pool elicited the greatest fraction of cross-reactive lymph node GC B cells that bound to both the monomeric RBD_Wu_ and monomeric RBD_XBB_ (34%), as compared to the RBD_Wu_-RBD_Wu_ + RBD_XBB_-RBD_XBB_ homodimer cocktail (25%) and RBD_Wu_-RBD_XBB_ heterodimer (19%) (p=0.024, Kruskall Wallis test) (Fig. 4A-D, Fig. S6 A-C). Notably, a variable subset of these cross-reactive B cells also bound to an RBD bait from SARS-CoV-1, a divergent sarbecovirus, (Fig. 4A-C,E, Fig. S7 A-C). Analysis of the fluorescence intensity associated with RBD monomer binding by the GC B cells elicited by the heterodimers and mixed homodimers indicated that the RBD_WX5_ pool elicited B-cells that were more consistently cross-reactive across individual mice (Fig. 4A-C, Fig. S6 A-C, Fig. 4F). Overall, these results suggest that incrementally bridging the genetic distance between the RBD_Wu_ and RBD_XBB_ using heterodimer components in the form of the RBD_WX5_ pool, rather than incorporating all the substitutions in a single heterodimer, elicits B cells and neutralizing antibodies with broader cross-reactivity.

**Fig. 4.**
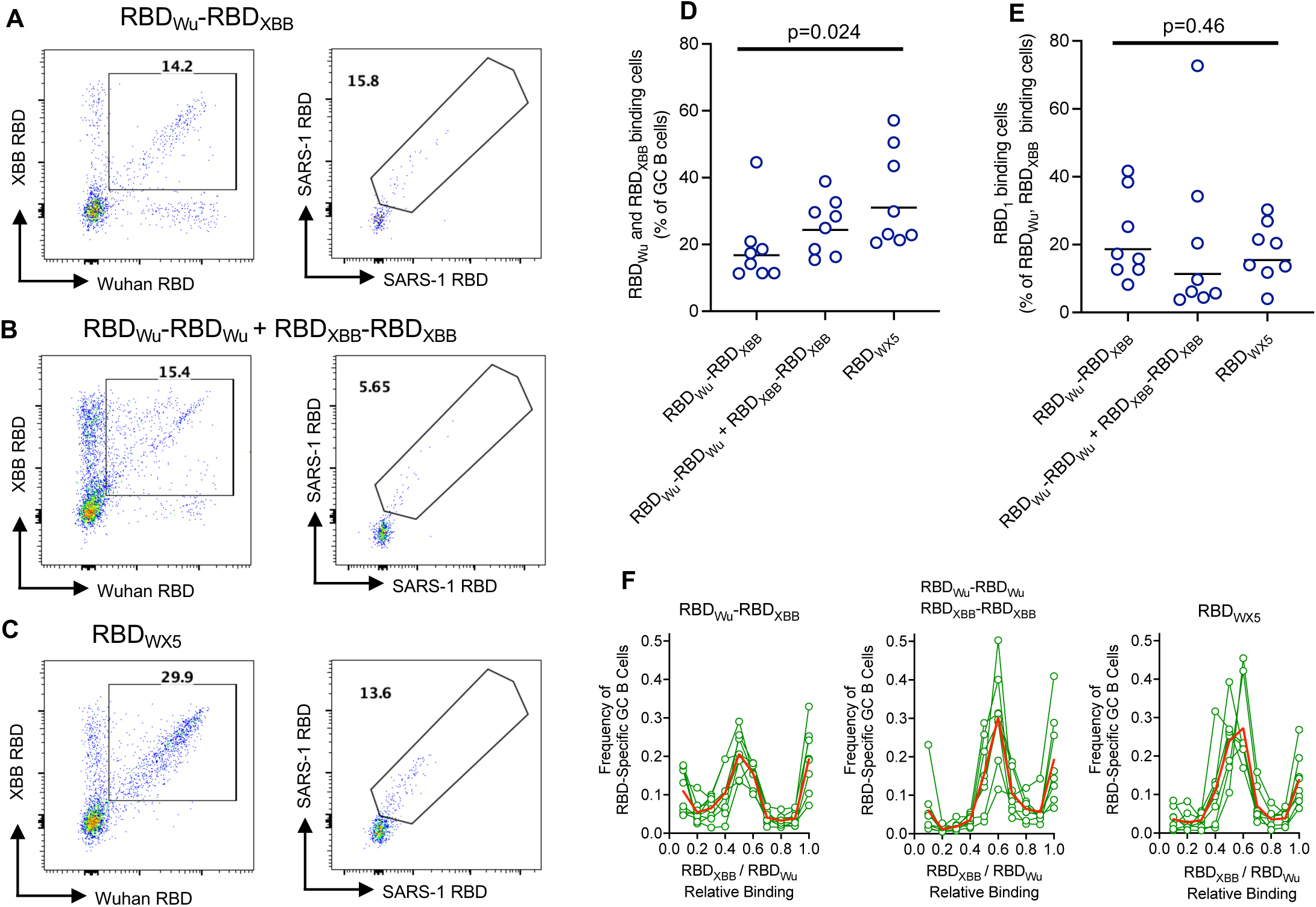
Cross-binding germinal center B cells elicited by overlapping RBD_WX5_ heterodimer pool. (A–C) Representative FACS plots of mouse lymph node germinal center B cells (B220^+^, CD4^−^, CD8^−^, NK1.1^−^, F4/80^−^, CD38^−^, and CD95^+^) binding to fluorophore-conjugated monomer RBD baits (RBD_Wu_, RBD_XBB_, RBD_S1_) for cells harvested on day 17 post-immunization. Bolded numbers represent the percentage of germinal center B cells that bind to RBD baits. (D) Percentage of GC B cells that bind both the RBD_Wu_ and RBD_XBB_ monomeric RBD baits. (E) GC B cells that bind all three monomeric RBD baits, expressed as a % of RBD_Wu_/RBD_XBB_ binding cells that also bind RBD_S1_. For (D) and (E) Lines represent group geometric mean, n = 8 mice per group. (F) Proportion of RBD-specific lymph node germinal center B cells (Y axis) that bind RBD_Wu_ and RBD_XBB_ monomer baits at the indicated relative fluorescence intensity ratios plotted on the X-axis. Graph title indicates immunogen group. Each green line represents one mouse, red lines indicate the group mean (n=8).

### Neutralization breadth of RBD_WX5_ heterodimer pool elicited antibodies

We next tested the ability of sera elicited by the RBD_Wu_ and RBD_XBB_ homodimers and heterodimers and the RBD_WX5_ pool to neutralize SARS-CoV-2 variants that were not included in the heterodimer pool (Fig. 3C). All immunogens that contained RBD_XBB_, including the RBD_WX5_ pool, generated high titer neutralizing antibodies (mean NT50 = 12,397 to 46,666) against SARS-CoV-2_BA.5_, an earlier variant that includes a subset of the substitutions present in SARS-CoV-2_XBB_ (Fig.5A. Fig. 3C). Conversely, a chimeric pseudotype containing the RBD from SARS-CoV-2_NJ_, an exceptionally divergent ‘cryptic’ variant recovered from New Jersey wastewater(47) (Fig. 3C) was resistant to neutralization by antibodies elicited by the individual or mixed RBD_Wu_ and RBD_XBB_ homodimers (mean NT50 = 20-137) but could be neutralized to some degree by sera from mice immunized with the RBD_Wu_-RBD_XBB_ heterodimer (mean NT50 = 727) and particularly the RBD_WX5_ pool (mean NT50 = 2,033) (Fig. 5B). We next tested mose sera for activity against a divergent sarbecovirus, SARS-CoV-1, that circulated in humans in 2002-2004. Neither the individual or pooled homodimers nor the the RBD_Wu_-RBD_XBB_ heterodimer elicited consistent neutralizing activity against SARS-CoV-1 (mean NT50 = 10-115) (Fig.5C). Only the RBD_WX5_ pool elicited neutralizing antibodies against SARS-CoV-1 in all six mice tested (mean NT50 = 455) (Fig. 5C). Clearly, sera from the RBD_WX5_ immunized mice had the greatest neutralization breadth, particularly against highly divergent sarbecoviruses SARS-CoV-2_NJ_ and SARS-CoV.

**Fig. 5.**
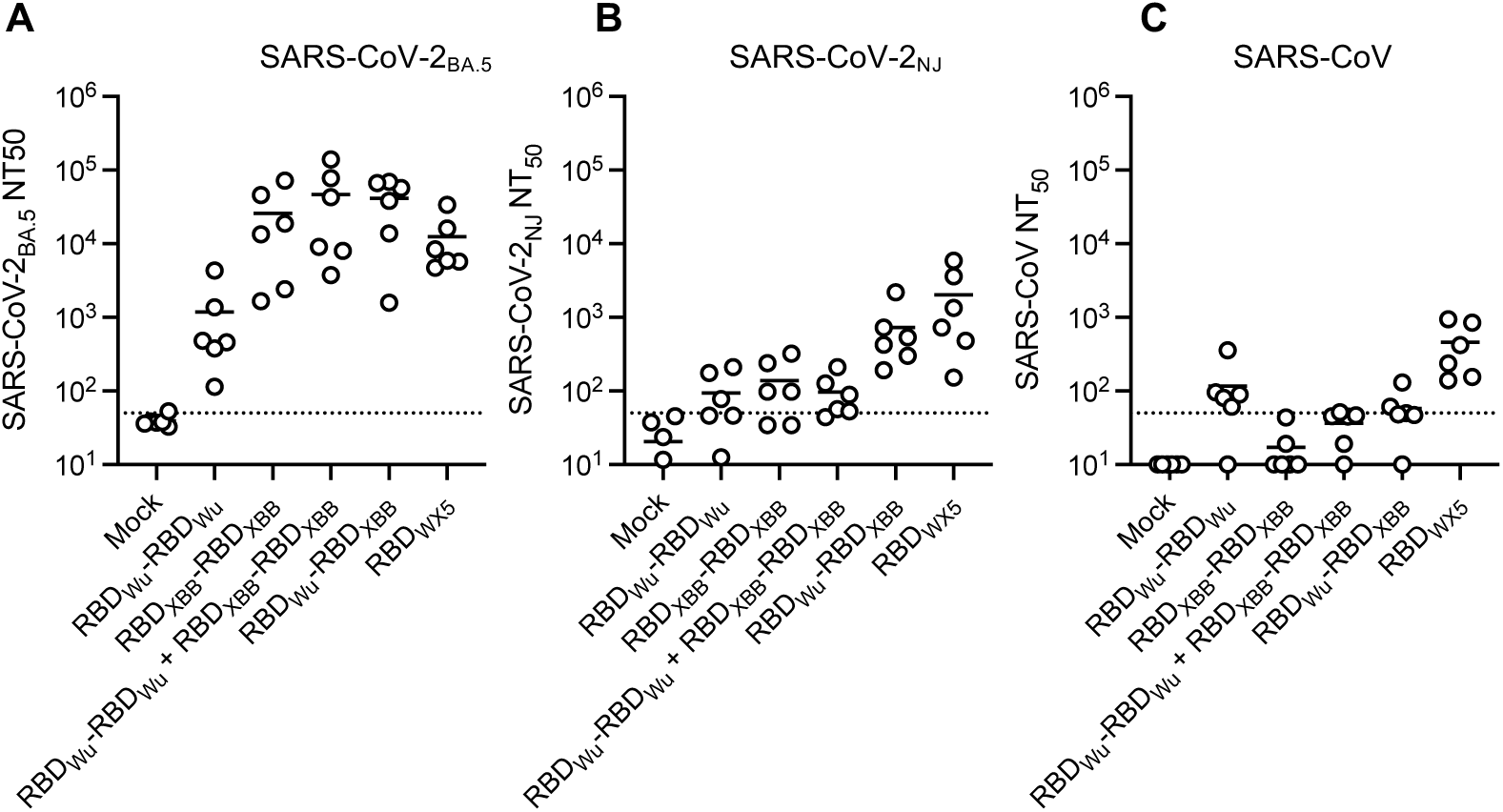
Neutralizing antibodies against heterologous SARS-CoV-2 variants and SARS-CoV-1 elicited by the RBD_WX5_ heterodimer pool. (A-C) Neutralizing titers (NT_50_) against SARS-CoV-2 pseudotypes; SARS-CoV-2_BA.5_ (A) and SARS-CoV-2_NJ_ (B) and SARS-CoV-1 (C) in mouse sera 12 weeks post-immunization with two doses of RBD homodimers or heterodimers, as indicated. Each symbol represents 1 mouse, lines represent group mean, n=5-6 mice per group. Dotted line indicates the lowest sera dilution tested (1:50).

We compared the serum neutralizing antibody response against SARS-CoV-2_Wu_ and SARS-CoV-2_XBB_ elicited by RBD homodimers and heterodimers to that elicited by mRNA-LNP vaccines encoding the full spike protein sequences of SARS-CoV-2_Wu_ and SARS-CoV-2_XBB_ (Fig. S8A-H). The SARS-CoV-2_Wu_ mRNA-LNP vaccine elicited little or no neutralizing activity against SARS-CoV-2_XBB_, and the SARS-CoV-2_XBB_ mRNA-LNP vaccine was similarly unable to elicit activity against SARS-CoV-2_Wu_ (Fig. S8A-B). Mixtures of spike mRNA-LNPs generated neutralizing antibodies against both SARS-CoV-2_Wu_ and SARS-CoV-2_XBB_ and the neutralizing titers elicited by mRNA-LNP vaccines and the RBD homodimers and heterodimers were similar. All immunogens were able to elicit neutralizing antibodies that plateaued at ∼7 weeks after immunization and were stable until at least week 12 (Fig. S8A-H).

### Efficacy of RBD heterodimer immunogens against ‘future’ SARS CoV-2 variants and divergent sarbecovirus

To test the durability of RBD dimer vaccines, and to avoid bias in the selection of heterologous SARS-CoV-2 strains with which to challenge experimental animals, we adopted an unusual ‘prospective’ experimental design at the time of RBD_WX5_ design and derivation (August 2023). Specifically, we undertook to immunize mice with the RBD_WX5_ heterodimer set and challenge them one year later with whatever SARS-CoV-2 strain was dominant at that time (i.e. in August 2024) (Fig. 6A). The immunogens included in this experiment included the homodimers RBD_Wu_-RBD_Wu,_ RBD_XBB_-RBD_XBB_, the corresponding heterodimer RBD_Wu_-RBD_XBB_, and RBD_WX5_ configured as a mixture of RBD monomers, or as a mixture of heterodimers as above. Mice were subjected to our standard 2 dose immunization schedule (Fig. 6A), bled at weeks 0, 3, 7, 12, 28 and 51 and sera was assessed for neutralizing activity against SARS-CoV-2_Wu_ and SARS-CoV-2_XBB_ for all weeks (Fig. 6B-F). In August 2024, one year after initiating this experiment, the predominant circulating SARS-CoV-2 strains were SARS-CoV-2_KP.2_ and SARS-CoV-2_KP.3.1_. Neutralizing titers against a hybrid of these two strains (SARS-CoV-2_KP.2/3_) were determined using mouse sera obtained at weeks 12, 28 and 51 post-immunization (Fig. 6B-F,

**Fig. 6.**
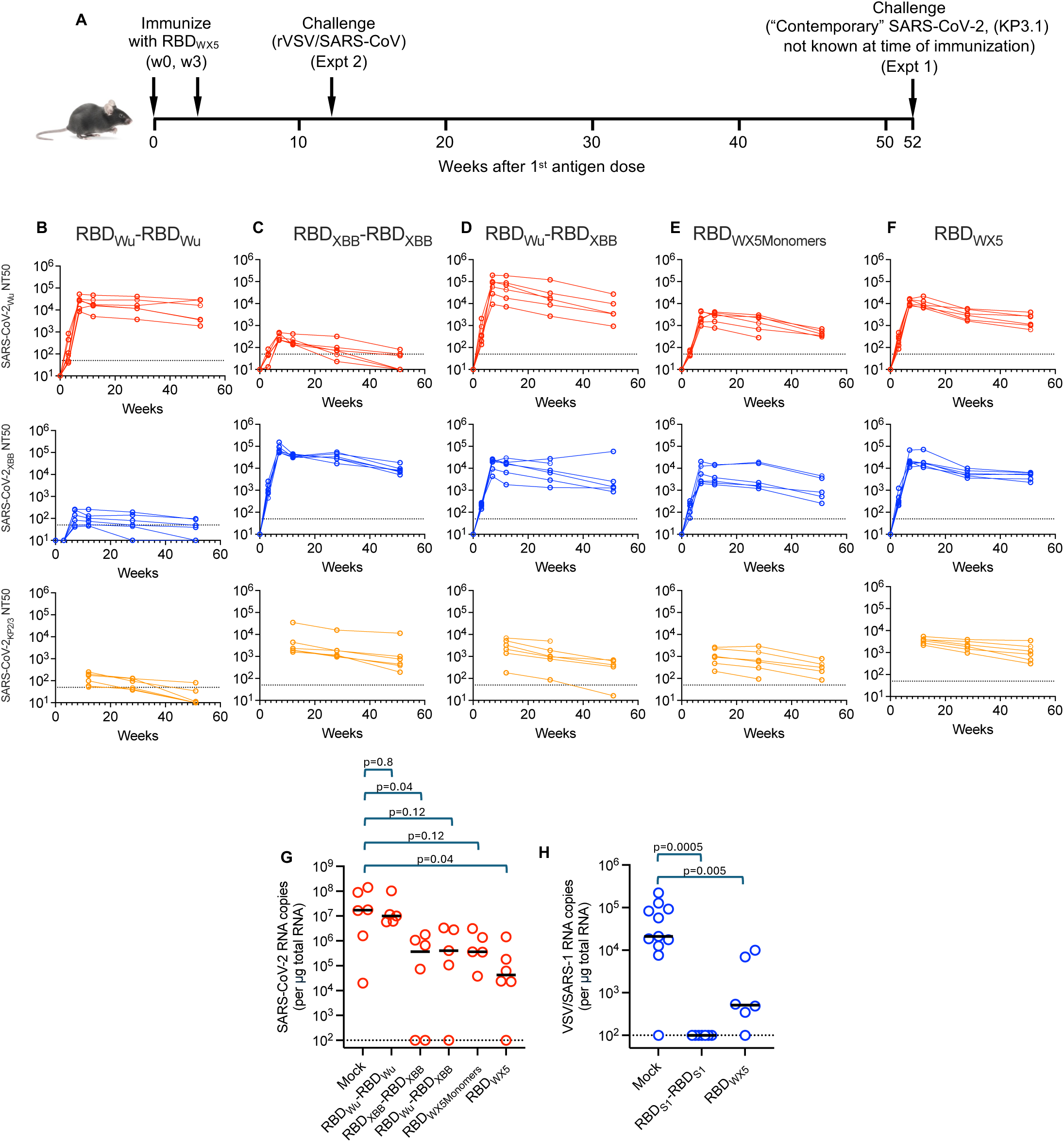
Neutralizing antibody persistence over one year and protection conferred by the synthetic RBD_WX5_ heterodimer pool. (A) Design and execution of an immunogen efficacy experiment to test the longevity of protection against SARS-CoV-2 variants that might emerge after immunization. Mice received two doses of immunogen, and were challenged one year later with whatever SARS-CoV-2 variant was prevalent one year after the design of the immunogen. (B-F) Neutralizing titers (NT_50_) over time (up to 1 year) against SARS-CoV-2_Wu_, SARS-CoV-2_XBB,_ SARS-CoV-2_KP2/3_ pseudotypes in mouse sera after immunization with two doses of the indicated immunogens. Graph title indicates immunogen, each line represents 1 mouse, n = 6 mice per group. Dotted line indicates the lowest sera dilution tested (1:50). (G) SARS-CoV-2_KP.3.1_ lung viral loads on day 3 after infection of K18-hACE2 mice, immunized one year previously with two doses of indicated immunogen. Each symbol represents mean of measurements from the left lung and right lung of each mouse, black line = group median. Dotted line indicates limit of detection. (H) Lung viral loads (rVSV/SARS-1 RNA copies per μg total RNA) on day 3 after infection of K18-hACE2 IFNAR(-/-) mice, immunized 12 weeks prior with two doses of indicated immunogen. Each symbol represents one mouse lung, each line represents group median. Dotted line indicates limit of detection.

Fig. S9A,B). As before (Fig. S4, Fig. S8), the RBD_Wu_-RBD_Wu,_ and RBD_XBB_-RBD_XBB_ homodimers elicited antibodies against the homologous strain and the RBD_XBB_-RBD_XBB_ homodimer also elicited neutralizing activity against a SARS-CoV-2_KP.2/3_ pseudotype (mean NT50 = 7,854 at 12 weeks). Notably, the natural rate of SARS-CoV-2 RBD evolution was slower than anticipated over the course of the year between antigen design/administration and challenge, resulting in circulating SARS-CoV-2 strains whose RBD differed at only three amino acid positions from RBD_XBB_ (F456L, Q493E and L455S) and thus was only modestly divergent from the RBD_XBB_ containing immunogens. The RBD_WX5_ heterodimer pool elicited neutralizing antibodies against SARS-CoV-2_Wu_, SARS-CoV-2_XBB_, and SARS-CoV-2_KP.2/3_ (mean NT50 = 10,688, 24,899 and 3,561 respectively at 12 weeks), values that were 2.9-fold to 4.8-fold higher than titers elicited by the same RBD_WX5_ mixture configured as monomers. This neutralizing activity was well maintained at 51 weeks, one week prior to challenge (mean NT50 = 2,021, 4,834 and 1,356 against SARS-CoV-2_Wu_, SARS-CoV-2_XBB_ and SARS-CoV-2_KP.2/3_ respectively) (Fig. 6B-F, Fig. S9A,B).

We obtained isolates of SARS-CoV-2_KP.3_ and SARS-CoV-2_KP.3.1_ (which have identical RBD sequences) and verified that these viruses replicated in K18-ACE2 mice (Fig. S9C). We challenged immunized mice at 52 weeks post-immunization to assess whether the neutralizing antibodies would confer protection against these “future” SARS-CoV-2 variants circulating one year after immunogen design and administration (Fig. 6A,G). Relative to unimmunized or RBD_Wu_-RBD_Wu_ immunized mice, those immunized with RBD_WX5_ had a median of >400-fold reduced lung viral burden following SARS-CoV-2_KP.3.1_ challenge (Fig.6G). Furthermore, RBD_WX5_ conferred better protection against SARS-CoV-2_KP.3.1_, compared to the RBD_XBB_-RBD_XBB_, RBD_Wu_-RBD_XBB_, and RBD_WX5 Monomers_ whose median reduction in viral burden was 47-, 42- and 47-fold compared to unimmunized mice. Together, these results suggest that RBD heterodimers, including RBD_WX5_, provide durable protection in mice against an evolved SARS-CoV-2 variant up to one year after a 2-dose immunization (Fig. 6G).

### RBD_WX5_ heterodimer pool confers protection against rVSV/SARS-CoV-1/GFP

To assess whether the SARS-CoV-2-based RBD_WX5_ confers protection against a divergent sarbecovirus, we developed a challenge model based on a recombinant VSV bearing the spike protein from SARS-CoV-1 (rVSV/SARS-CoV-1/GFP). This virus reproducibly infected and replicated in K18-hACE2 mice lacking functional type-I interferon receptors (IFNAR(-/-)) yielding ∼10^4^-10^5^ vRNA copies/µg total lung RNA, 72h after intranasal inoculation (Fig. S9D).

We immunized K18-hACE2/IFNAR(-/-) mice with a perfectly matched SARS-CoV-1 RBD homodimer (RBD_S1_-RBD_S1_), or the RBD_WX5_ pool, and then challenged them with rVSV/SARS-CoV/GFP. The matched RBD_S1_-RBD_S1_ homodimer provided apparently complete protection against rVSV/SARS-CoV-1/GFP infection. Notably, the RBD_WX5_ heterodimer pool that is highly divergent from SARS-CoV-1 (Fig. 3C) provided partial but substantial protection, reducing median rVSV/SARS-CoV-1/GFP lung viral burden by >40-fold (p=0.0005) compared to unimmunized control mice (Fig.6H).

## Discussion

Neutralization breadth, defined as the ability of a given antibody or collection of antibodies to neutralize a constellation of viral variants, might be achieved in mechanistically distinct ways. First, breadth could simply result from antibody targeting of epitopes that are conserved. Such conservation may be due to functional constraints, i.e high fitness costs associated with epitope variation. Alternatively, epitope conservation might also arise as consequence of the rareness of antibodies targeting a particular epitope. In such cases, viral populations remain ‘inexperienced’ to the effects of rare antibodies, and epitopes remain conserved due to the absence of selective pressure. A fundamentally different mechanism underlying neutralizing breadth arises when neutralizing antibodies attain high affinity, often through maturation, and as a result acquire an ability to tolerate variation in their epitope target(10, 39). Such affinity matured antibodies can neutralize a broader spectrum of variants than ancestral versions of the same antibody. In principle, each of these factors could contribute to neutralization breadth.

In practice, the serum neutralization breadth that we and others have found in individuals with multiple homologous or heterologous exposures to SARS-CoV-2 antigens was largely comprised of antibodies that target the RBD(12, 45, 48) and shows that repeated exposure to divergent SARS-CoV-2 spike antigens amplifies a cross-reactive RBD response rather than responses to more conserved regions of spike. While the need to bind ACE2 is expected to functionally constrain its sequence to a degree(18), the SARS-CoV-2 RBD is nevertheless intrinsically variable, due to the selection pressures placed on it by neutralizing antibodies. The SARS-CoV-2 neutralization breadth in human serum following repeated antigen exposure must therefore depend on antibodies that tolerate variation in their RBD epitopes, or target constrained RBD features. The near absence of neutralizing antibodies that survive depletion of serum with RBD and NTD proteins suggests that conservation of epitopes outside the RBD and NTD is because antibodies that bind to those regions are either ineffective or rare. Similarly, the greater conservation of other areas of the spike protein outside the RBD is likely due to the absence of selection pressure by neutralizing antibodies rather than due to functional constraints. Thus, despite the superficial attractiveness of conservation of other spike regions in considering vaccine designs, the human immune system is predisposed to generate cross-reactive RBD specific neutralizing antibodies, and clearly places evolutionary pressure on SARS-CoV-2 at this site.

In our initial comparison of RBD_Wu_-RBD_Wu_ and RBD_Beta+_-RBD_Beta+_ homodimer elicited antibodies, we found that RBD_Wu_-RBD_Wu_ elicited antibodies were less cross-reactive with RBD_Beta+_ than were RBD_Beta+_-RBD_Beta+_ elicited antibodies with RBD_Wu_. While the reasons for this phenomenon are unclear, the directional nature of this cross reactivity may be because the substitutions in RBD_Beta+_ relative RBD_Wu_ were chosen from substitutions that naturally occurred in RBD_Wu_ during the course of the COVID19 pandemic – i.e. they were naturally selected to reduce neutralizing antibody binding. The reverse is not the case; RBD_Wu_ amino acids were not naturally selected as escape variants of RBD_Beta+_. Thus, individual RBDs may differ in their propensity to bind individual antibodies elicited by their nearest neighbor in the pool.

Preferential stimulation of cross-reactive B cells by heteromultimeric immunogens via BCR crosslinking might be on the basis that those B cells (i) target portions of the heteromultimeric antigen that are conserved or (ii) tolerate variation in the targeted epitope(29). In our prior work we found that most SARS-CoV-2 neutralizing antibodies that had not undergone extensive affinity maturation over months could be escaped by single amino acid substitutions(5, 7, 39). Moreover, heterodimers with highly divergent RBD components generated separate sets of non-cross reactive neutralizing antibodies(38). Thus, in the design of the constituent dimers of the RBD_WX5_ heterodimer pool, we deliberately designed each heterodimer such that the number of amino acid substitutions for any given epitope in juxtaposed ‘nearest neighbor’ immunogens was minimized. Limiting the number of substitutions in each epitope of the heterodimer in this way should reduce the genetic barrier to stimulation of B cells that target the variable epitope, and thus lead to stimulation of greater numbers of B cells that are cross-reactive. Limiting the overall number of substitutions in a heterodimer also increases the amount of immunogen surface area that is fully conserved in the two components, and may favor stimulation of B cells that target those RBD epitopes.

Based on our findings with RBD_Wu_-RBD_Beta+_ immunized mice, RBD_WX5_ immunized mice should generate antibodies that bind to individual RBD components and cross react with their nearest neighbors in the RBD_WX5_ pool. However, it was unclear whether the generation of antibodies cross react with two antigenically distant RBDs would be enabled by the presence of intermediate RBD sequences. SARS-CoV-2_Wu_ and SARS-CoV-2_XBB_ neutralizing antibodies elicited by RBD_Wu_-RBD_Wu_ and RBD_XBB_-RBD_XBB_ homodimers or RBD_Wu_-RBD_XBB_ heterodimers, that represent the extremities of the RBD_WX5_ set, did not cross-react. In contrast, our depletion studies showed that there was cross reactivity among a subset of RBD_WX5_ elicited SARS-CoV-2_Wu_ and SARS-CoV-2_XBB_ neutralizing antibodies. We speculate that during affinity maturation, a naïve B-cell initially stimulated by (for example) a RBD_Wu_-RBD_Beta+_ heterodimer might generate variant BCRs that might further recognize RBD_Beta2+_ and therefore be efficiently stimulated by RBD_Beta+_-RBD_Beta2+_. Thus, in a stepwise manner, BCRs might be selected to recognize increasing numbers of heterodimers, with those recognizing the greatest numbers of heterodimers enjoying the greatest competitive advantage during the B cell response. Indeed, the finding that both SARS-CoV-2_Wu_ and SARS-CoV-2_XBB_ neutralizing antibodies were depleted by RBD_Beta3+_ (an intermediate in the RBD_WX5_ set) indicates that overlapping sets of cross-reactive antibodies are generated by RBD_WX5_.

Moreover, unlike any of the other homodimeric or heterodimeric RBD immunogens tested, the RBD_WX5_ pool generated antibodies that neutralized SARS-CoV, a highly divergent sarbecovirus, and conferred a substantial degree of protection against rVSV/SARS-CoV. Clearly the use of overlapping heterodimer pool generates collections of antibodies that have breadth, and some members of this antibody collection have cross-reactive properties that are rare or absent when only antigens at the extremities of the pool are included. Future work will need to examine in detail the properties and evolution of individual antibodies induced by heterodimer pools that attempt to guide the evolution of antibodies to a state that is broadly cross-reactive. Analysis of individual antibodies will also reveal to what extent the immunization approach described herein favors generation of antibodies that converge on particular target epitopes that are conserved among components of the mixture, to what extent the antibodies tolerate epitope variation and to what extent serum neutralization breadth is the consequence of individual antibody properties versus the behavior of collections of antibodies.

## Acknowledgements

This work was supported by grants from the National Institute of Allergy and Infectious Diseases P01AI165075 to (to PDB, TH, MCN, CMR and MRM), the Howard Hughes Medical Insititute (PDB and MCN) and the Stavros Niarchos Foundation Institute for Global Infectious Disease Research (MCN, CMR, TH and PDB). We thank the members of the Mount Sinai Pathogen Surveillance team for providing the infrastructure to identify, culture and provide rapid access to primary SARS-CoV-2 isolates.

## Materials and Methods

### Cell Lines

The cell lines 293T (CVCL_0063) and HT1080/ACE2.cl.14 (49) were cultured at 37°C and 5% CO_2_ in DMEM (ThermoFisher 11995065) containing 10% fetal bovine serum (Sigma F0926) and gentamicin (Gibco 15750060). Expi293 (ThermoFisher A14635) were cultured at 37°C and 8% CO_2_ in Expi293 Expression Medium (ThermoFisher A1435101) containing gentamicin (Gibco 15750060). Cell lines were periodically tested for retrovirus contamination using reverse transcriptase assays and mycoplasma contamination using DAPI staining.

### RBD_WX5_ heterodimer pool design

A series of four synthetic RBDs, termed RBD_Beta+_, RBD_Beta2+_, RBD_Beta3+_ and RBD_Beta4+_ that included subsets of the naturally occurring differences between SARS-CoV-2_Wu_ and SARS-CoV-2_XBB_, (the dominant variant circulating in August 2023) was designed using a structure and epitope guided approach. Starting with RBD_Wu_ and progressing through RBD_Beta+_, RBD_Beta2+_, RBD_Beta3+_ RBD_Beta4+_ and RBD_XBB_ each subsequent permutation in the series included four to five substitutions relative to the preceding RBD in the series, typically one from each of the four epitope classes, until the sequence of RBD_XBB_ was reached (excepting E484A and L368I). the series was termed “Wuhan-to-XBB x 5” (WX5). Double stranded DNA fragments encoding the RBD proteins of the RBD_WX5_ series were purchased GeneArt (ThermoFisher).

### RBD Beta Plus Series Sequences

#### Beta+

RFPNITNLCPFGEVFNATTFASVYAWNRKRISNCVADYSVLYNFASFSTFKCYGVSPTKLNDLCFTNVYADSFVIRGDEVRQIAPGQTGNIADYNYKLPDDFTGCVIAWNSNNLDSKVGGNYNYLYRLFRKSNLKPFERDISTEIYQAGSTPCNGVKGFNCYFPLQSYGFQPTYGVGYQPYRVVVLSFELLHAPATVCGPKKST

#### Beta2+

RFPNITNLCPFGEVFNATTFASVYAWNRKRISNCVADYSVLYNFASFSTFKCYGVSPTKLNDLCFTNVYADSFVIRGDEVSQIAPGQTGNIADYNYKLPDDFTGCVIAWNSNKLDSKVGGNYNYRYRLFRKSKLKPFERDISTEIYQAGSTPCNGVKGPNCYFPLQSYGFQPTYGVGYQPYRVVVLSFELLHAPATVCGPKKST

#### Beta3+

RFPNITNLCPFHEVFNATTFASVYAWNRKRISNCVADYSVLYNFASFSAFKCYGVSPTKLNDLCFTNVYADSFVIRGDEVSQIAPGQTGNIADYNYKLPDDFTGCVIAWNSNKLDSKPGGNYNYRYRLFRKSKLKPFERDISTEIYQAGSKPCNGVKGPNCYFPLRSYGFQPTYGVGYQPYRVVVLSFELLHAPATVCGPKKST

#### Beta4+

RFPNITNLCPFHEVFNATTFASVYAWNRKRISNCVADYSVLYNFASFFAFKCYGVSPTKLNDLCFTNVYADSFVIRGDEVSQIAPGQTGNIADYNYKLPDDFTGCVIAWNSNKLDSKPSGNYNYRYRLFRKSKLKPFERDISTEIYQAGNKPCNGVKGPNCYFPLRSYGFQPTYGVGHQPYRVVVLSFELLHAPATVCGPKKST

#### XBB

RFPNITNLCPFHEVFNATTFASVYAWNRKRISNCVADYSVLYNFAPFFAFKCYGVSPTKLNDLCFTNVYADSFVIRGNEVSQIAPGQTGNIADYNYKLPDDFTGCVIAWNSNKLDSKPSGNYNYLYRLFRKSKLKPFERDISTEIYQAGNKPCNGVKGPNCYSPLQSYGFRPTYGVGHQPYRVVVLSFELLHAPATVCGPKKST

### RBD and Spike Expression Plasmids

Plasmids encoding RBD_S1_, RBD_Wu_, RBD_Wu_-RBD_Wu_, and RBD_S1_-RBD_S1_ have been described previously(38). Sequences encoding the RBD of SARS-CoV-2_Wu_, SARS-CoV-2_XBB_ and components of the RBD_WX5_ series were amplified from previously described full-length spike plasmids(50, 51) or from synthetic double stranded DNA fragments (GeneArt). Plasmids expressing RBD monomers included an N-terminal secretory signal pSecTag2 (METDTLLLWVLLLWVPGSTGD) and a C-terminal 6xHisTag or an Avi-(GLNDIFEAQKIEWHE)-6xHisTag DNA fragments encoding the tagged proteins were inserted in the NcoI and XhoI sites of the mammalian expression plasmid pCAGGS. Plasmids expressing RBD dimers were constructed using three fragment ligation to insert (NcoI-pSecTag2-RBD-glycine-serine-linker(GGSGG)-NotI-RBD6xHisTag-XhoI) into the NcoI and XhoI sites of pCAGGS as described previously(38).

Plasmids encoding full-length SARS-CoV-1 and the SARS-CoV-2_Wu_ SARS-CoV-2_BA.5_ SARS-CoV-2_XBB_ variant spike proteins have been described previously (7, 10, 49–51). Plasmids encoding full-length SARS-CoV-2_Beta+_ and SARS-CoV-2_Beta3+_ spike proteins were constructed by replacing the RBD of SARS-CoV-2_Wu_ with the corresponding RBD_Beta+_ or RBD_Beta3+_ encoding sequences using an overlap extension PCR strategy followed by Gibson Assembly into a NheI/XbaI linearized pCR3.1 plasmid.

A plasmid expressing a chimeric SARS-CoV-2_Wu/KP2/3_ spike was made by introducing the point substitutions R346T/F456L/Q493E into a SARS-CoV-2_JN.1_, spike expression plasmid to generate the RBD_KP2/3_ sequence. An overlap extension PCR strategy was then used to replace the RBD encoding sequence of SARS-CoV-2_Wu_ spike with RBD_KP2/3_ encoding sequences, followed by Gibson assembly into an NheI/XbaI linearized pCR3.1 plasmid. A DNA sequence, encoding a SARS-CoV-2_NJ_ spike, a highly divergent sequence discovered during surveillance of New Jersey (NJ) wastewater, was commercially synthesized (ThermoFisher) and similarly inserted into an NheI/XbaI linearized pCR3.1 plasmid by Gibson assembly. Compared to the SARS-CoV-2_Wu_ spike sequence, the NJ wastewater spike contained the following substitutions and deletions: N334K, R346N, V367F, Y369S, del372, P384H, R403K, R408T, G413R, T415E, Y421F, N439K, N440T, K444N, V445N, G446K, Y449T, N450K, L452K, Y453F, L455F, F456V, N460R, K462Q, Y473F, S477N, T478R, G482T, del484, F486L, F490H, Q493L, S494T, G496N, N501S, Y508H, H519Q, K529R, N532D, E554D, D571E, T572N, Q580K, E583D, D614G, V615I, V622G, del638, S640P, V642G, Q675W, and P681S

A pre-fusion stabilized full length SARS-CoV-2_Wu_ spike encoding sequence was generated by mutating the furin cleavage site to G682SAS685 and introducing six prolines (P817,P892,P899,P942,P996,P997) by overlap extension PCR in an existing synthetic codon-optimized SARS-CoV-2_Wu_ spike sequence (49), lacking the C-terminal 19 codons, and encoding a C-terminal 6xHisTag. A pre-fusion stabilized version of SARS-CoV-2_XBB_ encoding the same modifications was generated by Gibson Assembly of several synthetic double stranded DNA fragments (GeneArt) into a XhoI/NotI linearized pCAGGS plasmid. Plasmids encoding the NTD domain of SARS-CoV-2_Wu_ and SARS-CoV-2_XBB_ were PCR amplified to incorporate an N-terminal secretory signal pSecTag2 and C-terminal 6xHisTag and inserted in the NcoI and XhoI sites of the mammalian expression plasmid pCAGGS.

### Protein Expression and Purification

NTD, RBD monomers and dimers, and pre-fusion stabilized spike proteins were produced by transfecting 75-600 million Expi293 cells with pCR3.1 or pCAGGS expression plasmids using the Expi293 Expression System (ThermoFisher A14635) according to manufacturer’s protocol. Culture supernatants were harvested four to six days post transfection, clarified by 0.22μm filtration (Millipore S2GPU05RE) and dialyzed into PBS using dialysis flasks (ThermoFisher 87762). Imidazole (1M) (Sigma I2399-100G) was added to a final concentration of 10 mM and the supernatant was batch bound to Ni-NTA Agarose beads (Qiagen 30210) overnight. Beads were collected by gravity flow into a column and washed with 10 column volumes of 50 mM NaH_2_PO_4_, 300 mM NaCl, 20 mM imidazole, pH 8.0. Proteins were eluted with 3 column volumes of 50 mM NaH_2_PO_4_, 300 mM NaCl, 250 mM imidazole, pH 8.0 and then dialyzed into PBS using Slide-A-Lyzer cassettes (ThermoFisher 66382). Protein quantity was calculated using the A_280nm_ (Nanodrop ThermoFisher) and the purity was visually assessed by separation on 4-12% Bis-Tris NuPage Gels (ThermoFisher NP0323BOX) followed by Gel Code Blue Staining (ThermoFisher 24590). Avi-Tagged RBD proteins were biotinylated using the BirA500 biotin-protein ligase standard reaction kit (Avidity EC 6.3.4.15) according to the manufacturer’s instructions, as previously described(13).

### Spike mRNA-LNP Formulation and Quantification

DNA templates, containing a T7 promoter region, 5’UTR, SARS-CoV-2 spike, 3’UTR, and poly-A tail, were linearized for *in vitro* transcription and mRNA transcript capping (5’-m7G Cap-1 structure) by T7 RNA polymerase and CleanCap Reagent AG (NEB E2080S). Uridine was substituted with the modified nucleoside pseudouridine (TriLink Biotechnologies, N-1019), at 5 mM, in the *in vitro* transcription reaction. DNase-I was then added to the reaction mixture, and the mRNA was purified by LiCl precipitation according to the manufacturer’s instructions (NEB E2080S). Next, 65 mg of mRNA were diluted in a total volume of 600 μL of 10 mM citrate buffer, and the following lipids were diluted in 300 μL of 100% ethanol at a 0.500: 0.015: 0.100: 0.385 molar ratio: ALC-0315 (Avanti Research, 890900O), ALC-0159 (Avanti Research, 880155P), DSPC (Avanti Research, 850365P), and cholesterol (plant) (Avanti Research, 700100P). A NanoAssemblr Ignite (Precision Nanosystems) was then used to mix the mRNA with the lipids at a flow rate of 8 μL/min. The resultant mRNA-LNP particles were dialyzed into 1X PBS + 10% sucrose overnight using Slide-A-Lyzer MINI dialysis devices with a 20kDa molecular weight cutoff (ThermoFisher, 88402). The Quant-iT RiboGreen RNA kit (ThermoFisher, R11490) was used to quantify the amount of mRNA in solution based on fluorescence emission intensity, relative to a ribosomal RNA standard curve. The mRNA-LNPs were treated with (or without) the detergent Triton X-100 (1%) to disrupt the lipid nanoparticles and release the encapsulated mRNA. The Quant-iT RiboGreen RNA reagent, a fluorescent nucleic acid stain, was then added to all samples. The difference in fluorescence signal between the detergent-treated (total mRNA) and detergent-untreated (non-encapsulated mRNA) samples was used to determine the concentration of the encapsulated mRNA-LNPs used for mouse immunizations.

### Mouse Immunizations

C57BL/6 (JAX Strain #000664) and K18-hACE2 mice were obtained from the Jackson Laboratory (JAX) and maintained in a standard ABSL1 or ABSL3 facility at the Rockefeller Comparative Biosciences Center. K18-hACE2 expressing IFNAR(-/-) mice were generated in-house and maintained in a standard ABSL2 facility. All mouse studies were performed in compliance with the Rockefeller University Institutional Animal Care and Use Committee (IACUC) according to animal protocol 24016-H. For analysis of neutralizing antibody responses, ten-week old C57BL/6 (JAX Strain #000664) and K18-hACE2 transgenic mice expressing the human ACE [B6. Cg-Tg (K18-ACE2)2Prlmn/J,034860] were immunized by intramuscular injection using a prime dose of 10μg of total protein in PBS (5 mg per leg) mixed with Enhanced Magic Mouse Adjuvant (Creative Diagnostics CDN-A001E) followed by a 10μg boost with the same protein 3 weeks later. For mRNA-LNP immunizations, mice were intramuscularly immunized with 1 mg of mRNA-LNP in a total volume of 50 mL of 1X PBS, followed by an identical ipsilateral booster dose 3 weeks post-prime. Mice were bled by the submandibular route at weeks 0, 3, 7, 12 weeks. For the one-year post immunization challenge experiment, mice were bled every 8 weeks post-week 12 until the time of viral challenge. For analysis of germinal center B cell resposnses, nine-week-old C57BL/6 (JAX Strain #000664) mice were immunized by footpad injection with 4μg of total protein in a 25-mL volume containing 8.3 mL of 2% Alhydrogel (Invivogen VAC-ALU-250) and 1X PBS. On day 17 post-immunization, draining popliteal lymph nodes were harvested and mechanically disrupted to create single-cell suspensions.

### Analysis of RBD-Binding properties of germinal center B Cells

For the RBD_Wu_-RBD_Beta+_ immunization experiments, biotinylated Avi-Tagged RBD_Wu_ (5μg/mL) and RBD_Beta+_ (5μg/mL) RBD baits were incubated with 1 μg/mL of either streptavidin-BUV661 (BD Biosciences 612979) or streptavidin-BV711 (BD Biosciences 563262) for 30 minutes.

Biotinylated RBD_XBB_ bait was conjugated to both streptavidin-APC (BioLegend 405207) and streptavidin-PE (BD Biosciences 554061). For the RBD_WX5_ immunization experiments, the biotinylated RBD_Wu_ bait was conjugated to streptavidin-BUV661, and the biotinylated RBD_XBB_ bait was conjugated to streptavidin-BV711. Biotinylated RBD_S1_ RBD bait was conjugated to both streptavidin-APC and streptavidin-PE. The RBD-streptavidin conjugates were then pooled, and a 1:200 dilution of Mouse BD Fc Block (BD Biosciences 553142) and a 1:500 dilution of Zombie NIR (BioLegend 423106) were added to the mixture. Cells were stained with 200 μL of the RBD-streptavidin complexes for 30 minutes, centrifuged at 350 g for 5 minutes, and stained with an antibody cocktail containing a 1:200 dilution of the following antibodies for 20 minutes: 1) anti-CD4-APC-eFluor780 (ThermoFisher 47004282), 2) anti-CD8a-APC-eFluor780 (ThermoFisher 47008182), 3) anti-NK1.1-APC-eFluor780 (ThermoFisher 47594182), 4) anti-F4/80-APC-eFluor780 (ThermoFisher 47480182), 5) anti-CD38-AlexaFluor 700 (ThermoFisher 56038182), 6) anti-CD95-BUV563 (BD Biosciences 741292), 7) anti-CD45R/B220-BUV805 (BD Biosciences 748867), 8) anti-igD-BV786 (BD Biosciences, 563618), 9) anti-CD138-BV421 (BioLegend, 142507), and 10) anti-GL7-FITC (BioLegend, 144604). Finally, cells were washed with 200 μL of 1X PBS, filtered through a 40 μM strainer (Fisher Scientific 08-771-1), and fluorescence profiles acquired on a BD FACSymphony cytometer (BD Biosciences). Germinal center (GC) B cells were defined as B220^+^, CD4^−^, CD8^−^, NK1.1^−^, F4/80^−^, CD38^−^, CD95^+^, and the RBD binding cells in this fraction were enumerated.

To calculate the relative binding propensities of the lymph node GC B cells to the fluorophore-conjugated RBD_Wuhan_ (BUV661) and RBD_Beta+_ (BV711) baits, the fluorescence intensities of RBD_Wuhan_ and RBD_Beta+_ -binding GC B cells were exported from the FlowJo software (10.9.0). For each of these RBD-specific cells, the RBD_Beta+_-BV711 fluorescence intensity was divided by the sum of the of RBD_Beta+_-BV711 fluorescence intensity and the RBD_Wuhan_-BUV661 fluorescence intensity to obtain a binding propensity ratio between 0 and 1. On the basis of these ratios, the cells were then grouped into 0.1-increment “bins”, and raw cell counts in each “bin” were divided by the total number of acquired RBD_Wuhan_ and RBD_Beta+_ -binding GC B cells to obtain relative frequencies. Cells with negative fluorescence intensity values were excluded from these calculations.

### Pseudotyped HIV-1 reporter virus production

Stocks of SARS-CoV-1 and SARS-CoV-2 spike pseudotyped human immunodeficiency virus (HIV-1)-NanoLuc were produced in 293T cells using polyethylenimine (Polysciences Cat# 23966) co-transfection of pHIV-1_NL4-3_ ΔEnv-NanoLuc and a pCR3.1 derived plasmid expressing full length-spike as described previously (49). Supernatants were harvested 48 hr post transfection, 0.22 μm filtered (Millipore S2GPU05RE), aliquoted and store at -80°C. Virus titers were determined by serial dilution and infection of HT1080/ACE2.cl14, NanoLuc luciferase activity 48 hr post infection was measured using the Nano-Glo Luciferase Assay System (Promega N1150) and a Glomax Navigator luminometer (Promega) cells as described previously (49).

### Neutralization Assays

Mouse serum and human plasma were tested for neutralizing activity against SARS-CoV-1 and SARS-CoV-2 and its variants using the SARS-CoV-2 spike-pseudotyped HIV-1 assay previously described (49). Mouse and human samples were heat-inactivated at 55°C for 30 min and then five-fold serial diluted using a Pipetmax (Gilson GFAM0072). Diluted serum or plasma was incubated with spike pseudotyped HIV-1 reporter virus for 1 h at 37 °C and then transferred onto HT1080/ACE2.cl14 cells. 48 hrs post infection the cells were washed with PBS, lysed in Luciferase Cell Culture Lysis reagent (Promega E1531), and luciferase activity was measured using the Nano-Glo Luciferase Assay System (Promega N1150) and a ClariostarPlus Microplate Reader (BMG LabTech). NT_50_ values were calculated the using four-parameter non-linear regression of normalized relative luminescence units (RLUs) plotted in GraphPad Prism as described previously (49).

### Neutralization Depletion Assays

Neutralization depletion assays were performed as described previously with slight modifications(38). For mouse serum samples, His-Tag Dynabeads (ThermoFisher 10103D) were pre-loaded with RBD-6xHis at a ratio of 10 µg RBD-6xHis per 10 µl His-Tag Dynabeads in PBS for 30 min at 4°C with end over end rotation. The beads were washed with PBS and aliquoted into 96 well plates containing 4 µL of mouse serum per well. For human plasma samples, His-Tag Dynabeads were pre-loaded with pre-fusion stabilized spike-6XHis, RBD-6xHis, or NTD-6xHis at a ratio of 10 µg 6XHis Protein per 19 µl His-Tag Dynabeads in PBS for 30 min at 4°C with end over end rotation. The beads were washed with PBS and aliquoted into 96 well plates containing 10 µL of human plasma per well. After 60 min incubation at 4°C with shaking, the beads were removed by magnetic separation. The supernatant was transferred to a fresh 96 well plate and screened for remaining neutralizing activity using the spike-pseudotyped HIV-1 assay.

### Human Plasma Samples

Plasma samples were obtained from human volunteers who were described in detail in previous studies. (7, 40, 43, 52)

### SARS-CoV-2 isolates

The Mount Sinai Pathogen Surveillance Program leverages residual clinical biospecimen that tested positive for SARS-CoV-2 to track the spread and evolution of disease-causing viral variants. The study protocol was IRB-approved (STUDY-13-00981). Briefly, viral RNA was extracted from viral transport media and complete viral genomes were generated as described (PMID 32471856, 40774230). Specimen were selected for viral culture on Vero-E6 cells expressing transmembrane protease serine 2 (TMPRSS2) based on the viral genome sequence information as previously described (PMID 37270625). The viral isolates hCoV-19/USA/NY-MSHSPSP-PV111989/2024 and hCoV-19/USA/NY-MSHSPSP-PV107407/2024 are representatives of clade 24E (KP.3.1.1) which circulated in NYC in the summer of 2024.

### SARS-CoV-2 Challenge Experiments

K18-hACE2 mice were infected intranasally with 20,000 PFU of SARS-CoV-2_KP3.1_ in a 25 μL volume. At day 3 post-infection, the right and left lungs were harvested from each mouse for RNA extraction and RT-qPCR. The lungs were first homogenized in Trizol LS reagent, and total RNA was extracted by phase separation using chloroform. RNA in the aqueous phase was precipitated with isopropanol, and pelleted RNA was washed with 75% ethanol. The RNA was dissolved in 100 μL nuclease-free water, and the number of viral genomes per microgram of lung was measured by qRT-PCR using 1-Step Kit, PowerSYBR Green RNA-to-CT (ThermoFisher 4389986) and StepOne Plus Real-Time PCR system (Applied Biosystems). The primers used were 2019-nCoV_N1-F: 5′-GACCCCAAAATCAGCGAAAT-3′ and 2019-nCoV_ N1-R: 5′-TCTGGTTACTGCCAGTTGAATCTG-3′, targeting RNA sequences that encode the nucleocapsid protein of SARS-CoV-2_KP3.1_. A standard, 2019-nCoV_N_Positive Control 10006625 was obtained from IDT.

### rVSV/SARS-CoV-1/GFP Production and Challenge

rVSV/SARS-CoV-1/GFP virus was produced using a previously described rescue protocol for a replication-competent rVSV/SARS-CoV-2 chimera (49). In rVSV/SARS-CoV-1/GFP the SARS-CoV-1 spike replaces the native VSV-G. K18-hACE2/IFNAR(-/-) mice were infected intranasally with 10^6^ IU of rVSV/SARS-CoV-1/GFP in a 25 μL volume. At day 3 post-infection, the left lung was harvested from each mouse and subsequently homogenized for downstream RNA extraction and qRT-PCR, as described above. The primers used were 5′-CTCTGCCGACTTGGCACAAC-3’ and 5′-TTCAAACCATCCGAGCCATTCG-3’, targeting RNA sequences that encode the nucleoprotein (N) of VSV. A reference plasmid, containing VSV N, was used as a standard to interpolate the number of viral RNA copies per microgram of input lung RNA in each reaction.

---

**Fig. S1.**
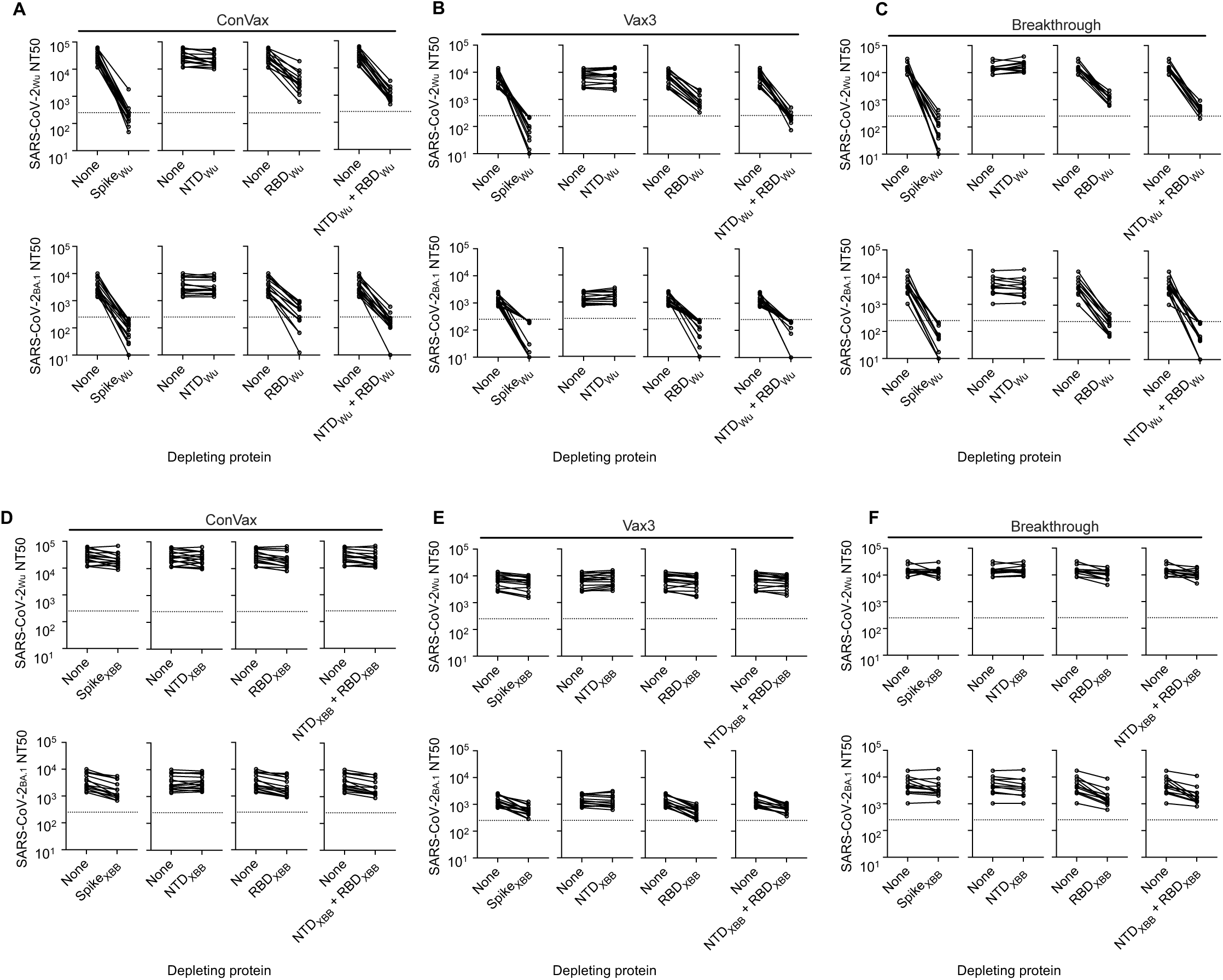
Neutralizing Antibodies are Largely Elicited by the RBD of SARS-CoV-2. (A-C) Neutralizing antibody titers (NT_50_) against SARS-CoV-2_Wu_ or SARS-CoV-2_BA.1_ pseudotype variants following mock depletion of patient plasma (None) or depletion by the indicated SARS-CoV2_Wu_ Spike(S), NTD(N), RBD(R), or combined NTD and RBD(N+R) proteins. (D-F) Neutralizing antibody titers (NT_50_) against SARS-CoV-2_Wu_ or SARS-CoV-2_BA.1_ pseudotype variants following mock depletion of patient plasma (None) or depletion by the indicated SARS-CoV2_XBB_ Spike(S), NTD(N), RBD(R), or combined NTD and RBD(N+R) protein. Graph title indicates patient group, each line represents 1 participant, ConVax n=15, Vax3 n=15, Breakthrough n=13. Dotted line indicates the lowest serum dilution tested (1:250).

**Fig. S2.**
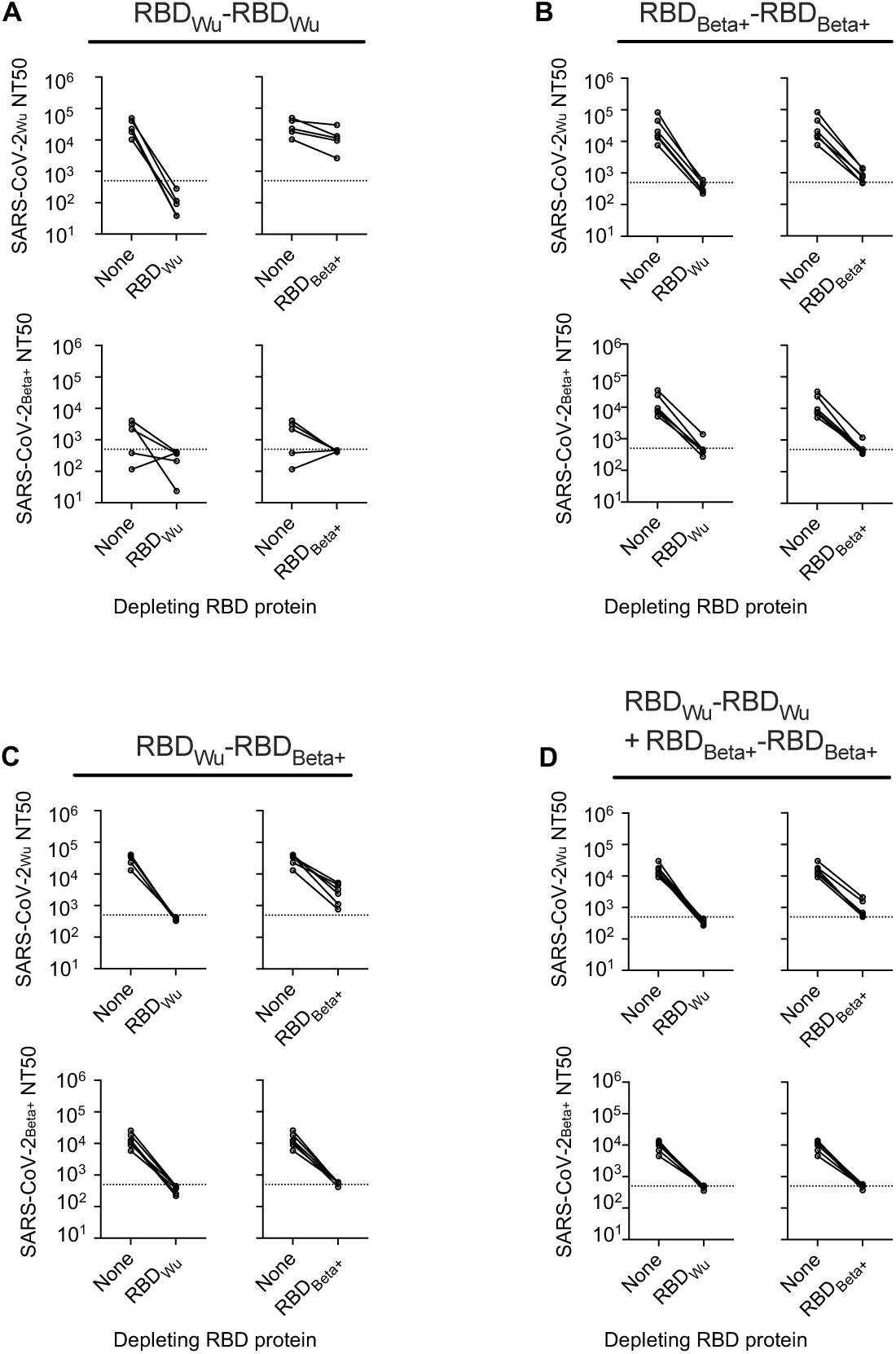
Cross-reactive neutralizing antibodies induced in escape homodimer and heterodimer immunized mice. (A-D) Neutralizing titers (NT_50_) against SARS-CoV-2 variant pseudotypes in 12 week post-immunization mouse sera following mock depletion (None) or depletion by the indicated RBD protein. Graph title indicates the immunogen used, each line represents1 mouse, n = 4-6 mice per group. Dotted line indicates the lowest serum dilution tested (1:500).

**Fig. S3.**
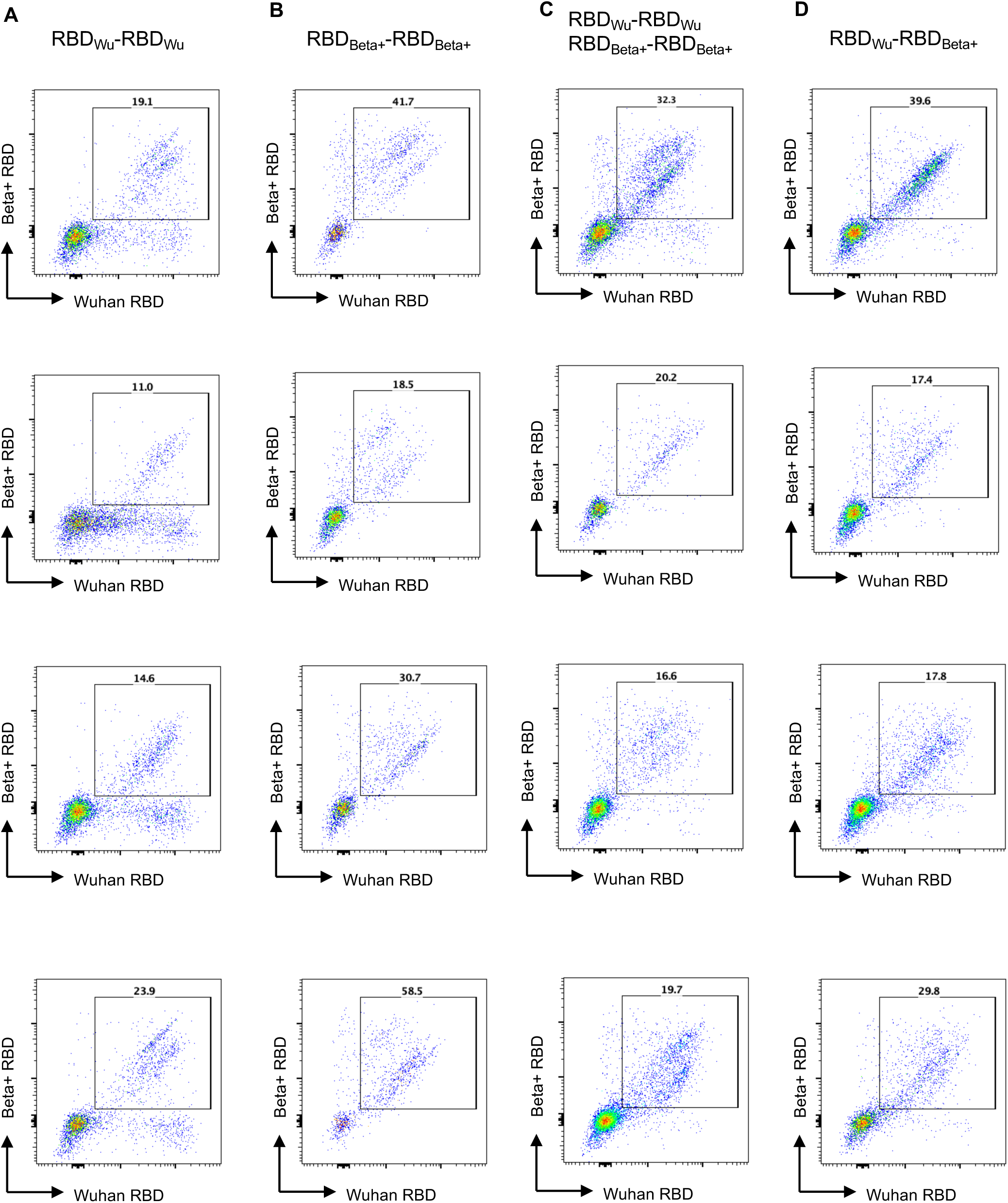
Cross-reactive B cells induced in RBD homodimer and heterodimer immunized mice. (A-D) FACS plots showing mouse lymph node germinal center B cells binding to fluorophore-conjugated RBD_Wu_ and RBD_Beta+_ baits 17 days after footpad immunization with the individual RBD_Wu_-RBD_Wu_ (A) or RBD_Beta+_-RBD_Beta+_ (B) homodimers, a mixture of RBD_Wu_-RBD_Wu_ and RBD_Beta+_-RBD_Beta+_ homodimers (C) or a RBD_Wu_-RBD_Beta+_ heterodimer (D). Numbers indicate the percentage of cells within the gate (that represents binding to both RBD_Wu_ and RBD_Beta+_ monomer baits). Graph title indicates immunogen group. Each plot represents 1 mouse, n=4 mice per group.

**Fig. S4.**
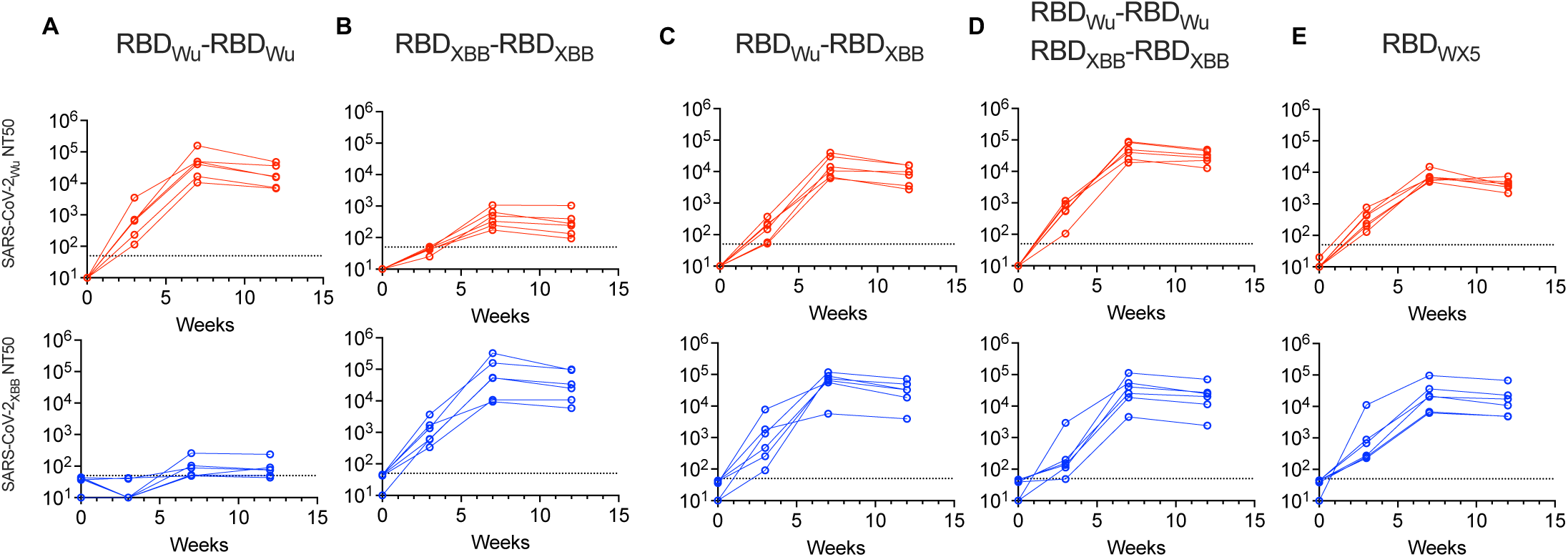
Neutralizing antibodies elicited by RBD dimers over 12 weeks. (A-E). Neutralizing titers (NT_50_) over time against SARS-CoV-2_Wu_ and SARS-CoV-2_XBB_ pseudotypes in mouse sera after immunization with two doses (week 0 and week 3) of the indicated immunogens. Graph title indicates immunogen, each line represents 1 mouse, n=6 mice per group. Dotted line indicates the lowest sera dilution tested (1:50).

**Fig. S5.**
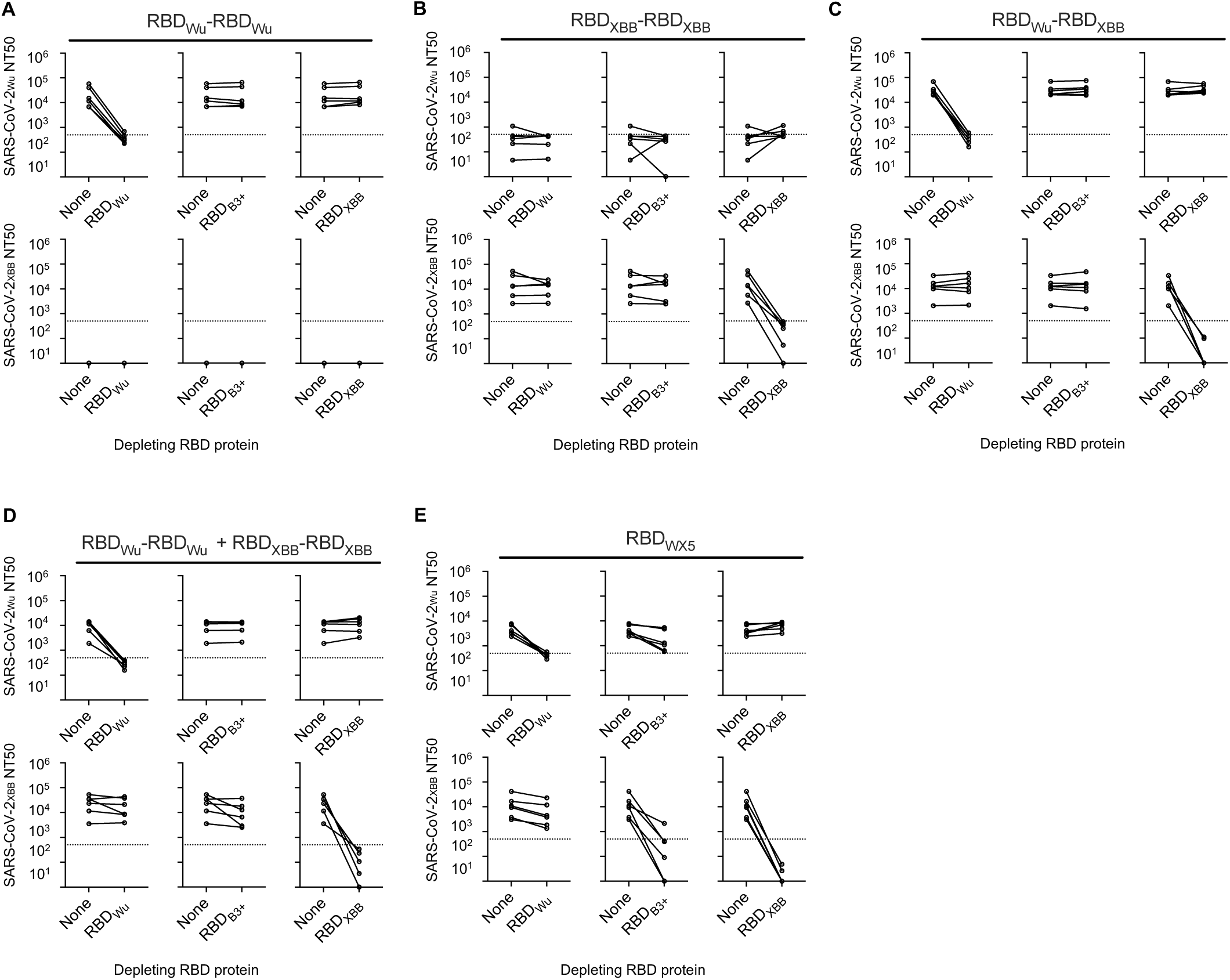
Cross-reactivity of neutralizing antibodies elicited by RBD homodimers, heterodimers and the tiled heterodimer series. (A-G) Neutralizing titers (NT_50_) against SARS-CoV-2_Wu_ and SARS-CoV-2_XBB_ pseudotypes in 12 week post-immunization mouse sera following mock depletion (None) or depletion by the indicated RBD proteins. Graph title indicates immunogen, each line represents 1 mouse, n = 4-6 mice per group. Dotted line indicates the lowest serum dilution tested (1:500).

**Fig. S6.**
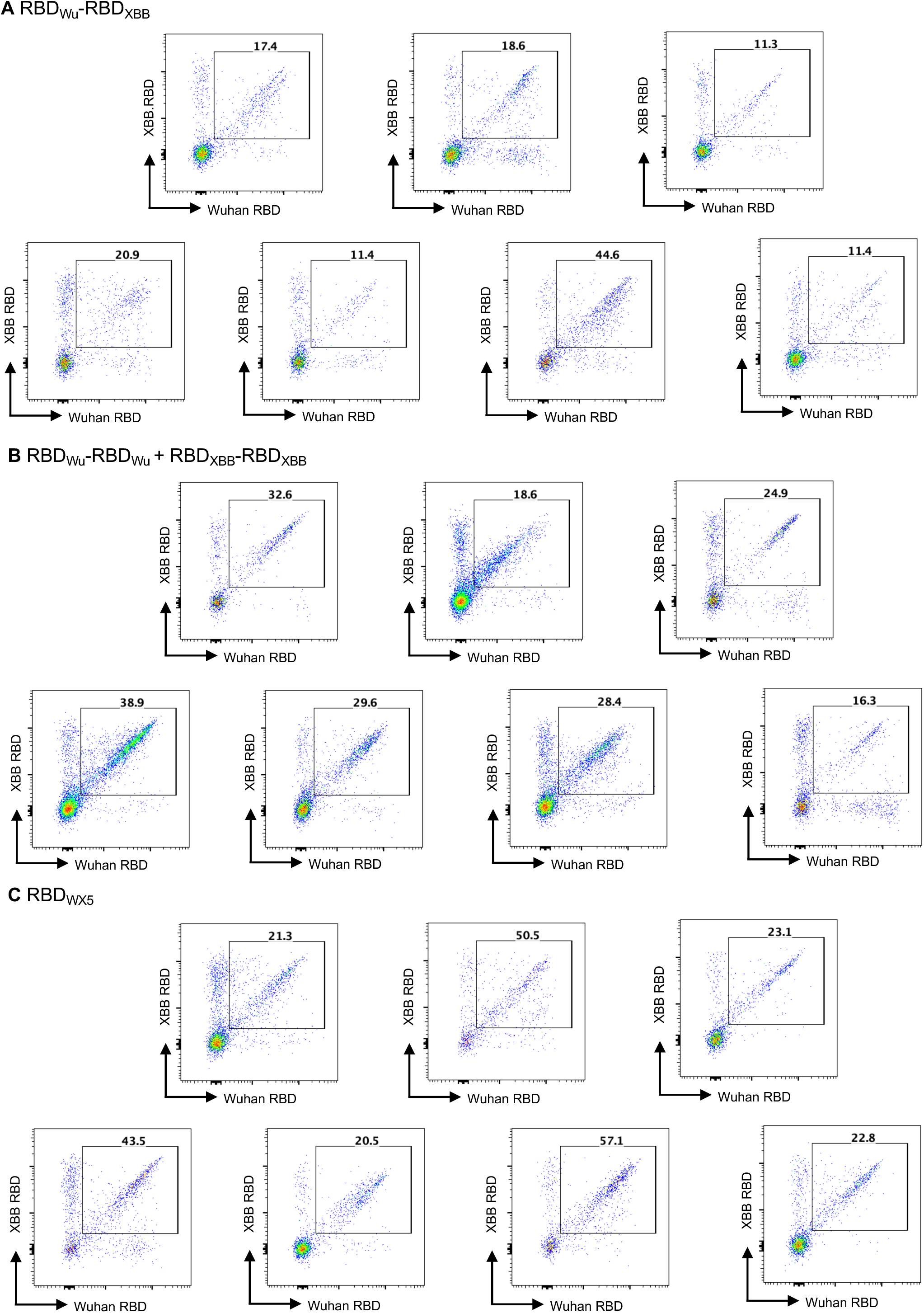
RBDWu/RBDXBB Cross-binding germinal center B cells elicited by a single RBD_Wu_-RBD_XBB_ heterodimer, mixed RBD_Wu_-RBD_Wu_/RBD_XBB_-RBD_XBB_ homodimers, and the RBD_WX5_ heterodimer pool. (A–C) FACS plots of mouse lymph node germinal center B cells (B220^+^, CD4^−^, CD8^−^, NK1.1^−^, F4/80^−^, CD38^−^, and CD95^+^) binding to fluorophore-conjugated monomer RBD baits (RBD_Wu_, RBD_XBB_) for cells harvested on day 17 post-immunization. Bolded numbers represent the percentage of germinal center B cells that bind to both monomeric RBD_Wu_ and RBD_XBB_ baits.

**Fig. S7.**
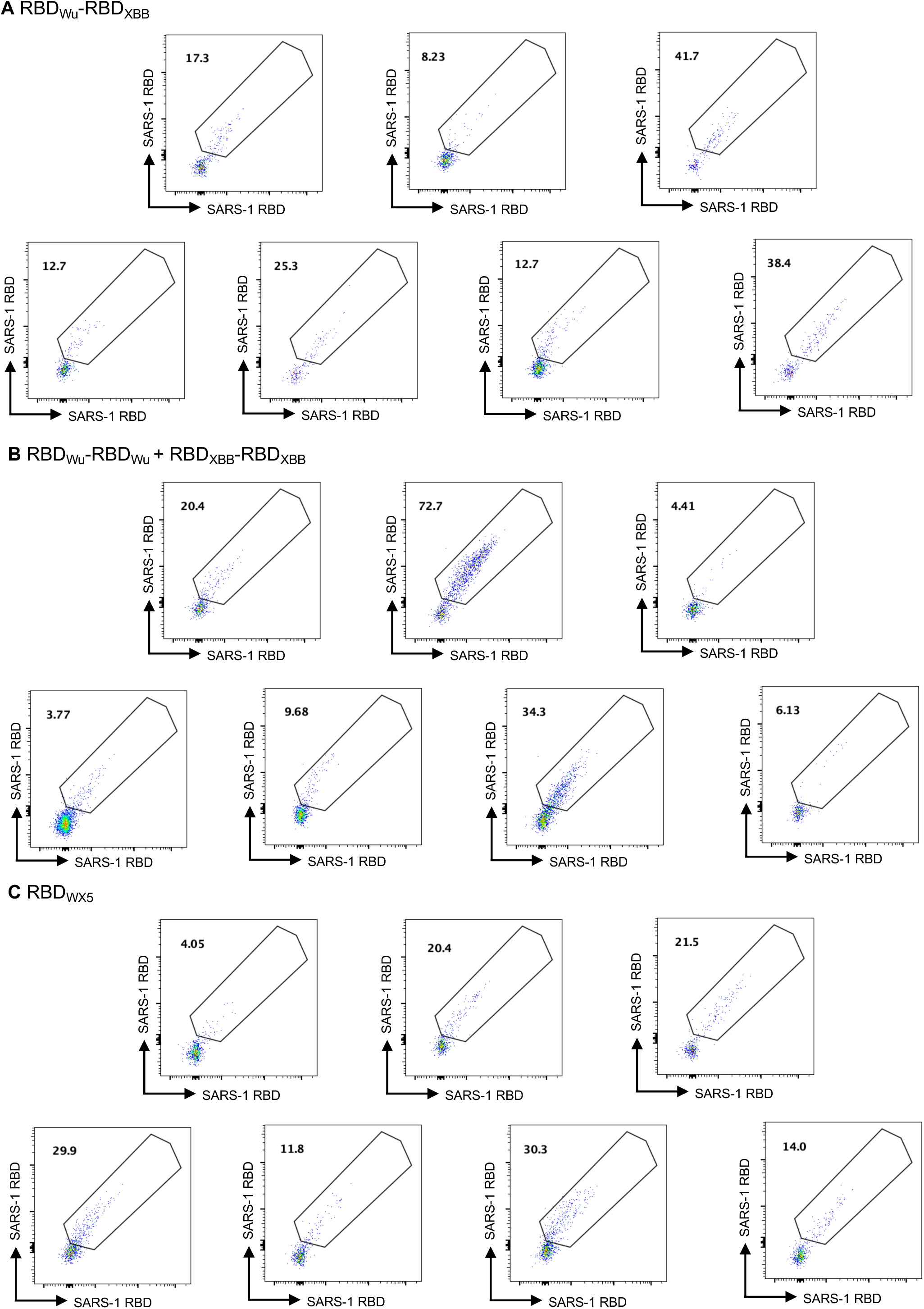
RBDWu/RBDXBB/RBDS1 Cross binding germinal center B cells elicited a single RBD_Wu_-RBD_XBB_ heterodimer, mixed RBD_Wu_-RBD_Wu_/RBD_XBB_-RBD_XBB_ homodimers, and the RBD_WX5_ heterodimer pool. (A–C) FACS plots of Fluorophore conjugated RBD_S1_ binding to mouse lymph node germinal center B cells (B220^+^, CD4^−^, CD8^−^, NK1.1^−^, F4/80^−^, CD38^−^, and CD95^+^), gated on cells that bound to both RBD_Wu_, and RBD_XBB_. Bolded numbers represent the percentage of RBD_Wu_/RBD_XBB_ cross reactive B cells that also bind to RBD_S1_.

**Fig. S8.**
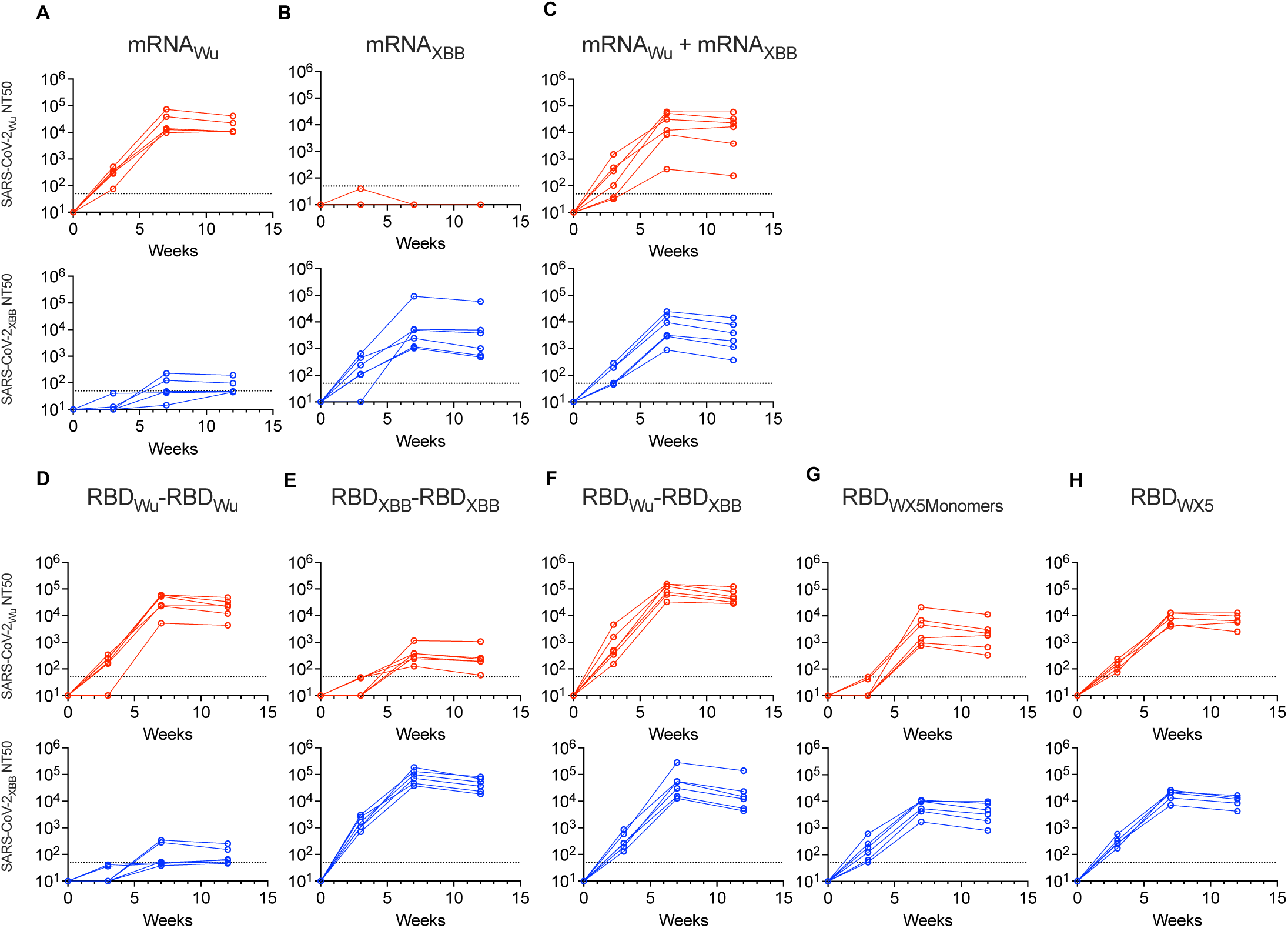
Comparison of neutralizing antibodies elicited by mRNA-LNP or RBD dimer protein immunogens over time. (A-H) Neutralizing titers (NT_50_) over time against SARS-CoV-2_Wu_ and SARS-CoV-2_XBB_ pseudotypes in mouse sera after immunization with two doses of the indicated immunogens. Graph title indicates immunogen, each line represents 1 mouse, n = 5-6 mice per group. Dotted line indicates the lowest sera dilution tested (1:50).

**Fig. S9.**
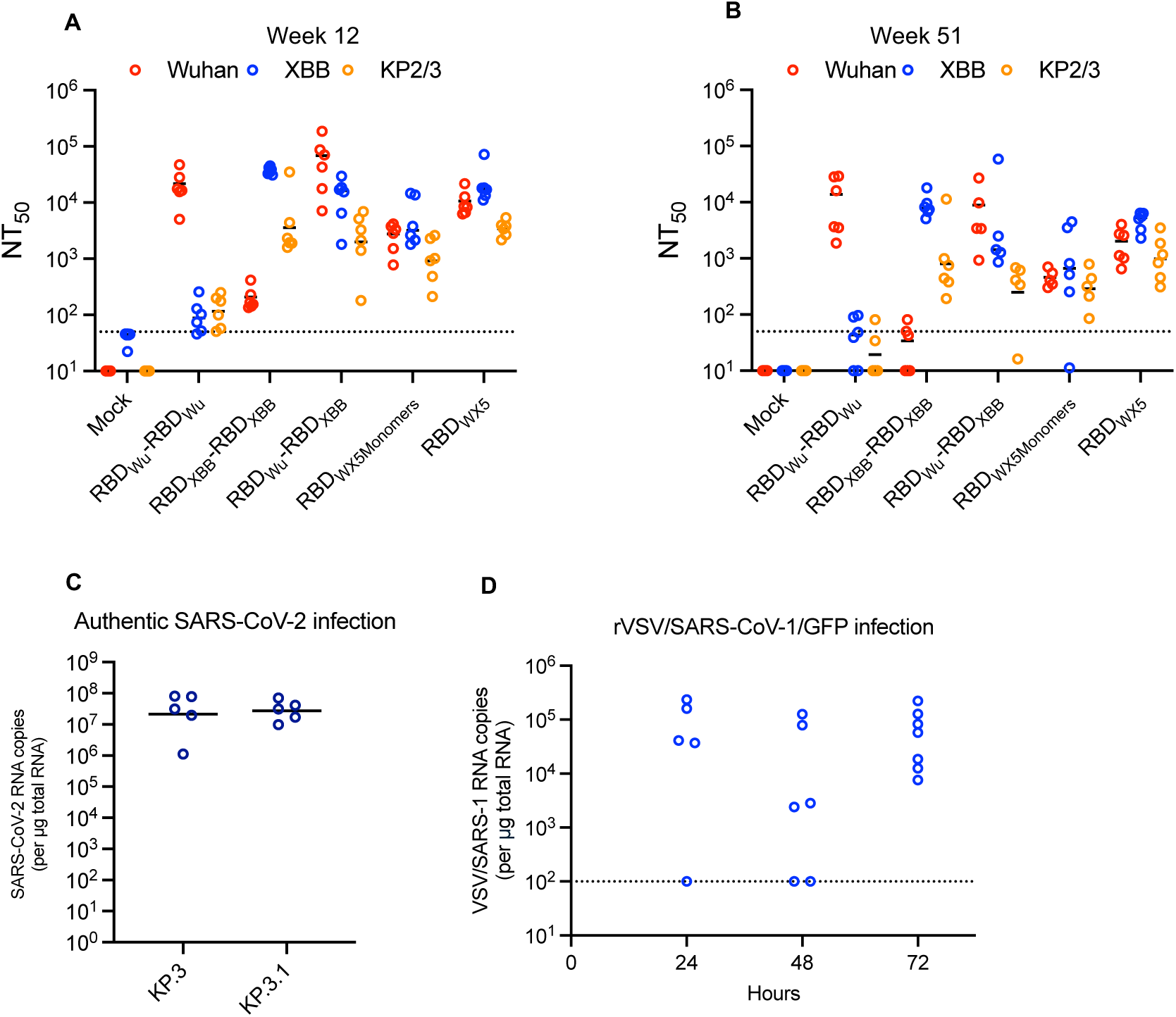
Neutralizing antibodies and virus challenge stocks for evaluation of tiled RBD heterodimer immunogen. (A,B) Comparison of neutralizing titers (NT_50_) against SARS-CoV-2 variant pseudotypes in mouse sera 12 and 51 weeks post-immunization with two doses of indicated immunogen. Each symbol represents 1 mouse, lines = group mean, n = 5-6 mice per group. Dotted line indicates the lowest sera dilution tested (1:50). (C) Validation of SARS-CoV-2 challenge virus stocks, KP.3 and KP.3.1. Lung viral loads (SARS-CoV-2 RNA copies per μg total RNA) on day 3 after infection of K18-hACE2 mice. Each symbol represents 1 mouse, lines = group geometric mean, n=5 mice per group (D) Lung viral loads (rVSV/SARS-1 RNA copies per μg total lung RNA) on each of the indicated hours after infection of K18-hACE2 IFNAR(-/-) mice. Each symbol represents 1 mouse, dotted line=limit of detection.

## References

1. E. Adjei-Gati et al., Harnessing conserved epitopes and multivalent antigen strategies for vaccine design: Lessons learned and opportunities. Hum Vaccin Immunother 22, 2652680 (2026).

2. W. J. Moss, D. E. Griffin, What’s going on with measles? J Virol 98, e0075824 (2024).

3. P. J. Klasse, R. W. Sanders, A. B. Ward, I. A. Wilson, J. P. Moore, The HIV-1 envelope glycoprotein: structure, function and interactions with neutralizing antibodies. Nat Rev Microbiol 23, 734–752 (2025).

4. R. Uraki, B. Korber, M. S. Diamond, Y. Kawaoka, SARS-CoV-2 variants: biology, pathogenicity, immunity and control. Nat Rev Microbiol 24, 8–28 (2026).

5. Y. Weisblum et al., Escape from neutralizing antibodies by SARS-CoV-2 spike protein variants. Elife 9 (2020).

6. A. Baum et al., Antibody cocktail to SARS-CoV-2 spike protein prevents rapid mutational escape seen with individual antibodies. Science 369, 1014–1018 (2020).

7. Z. Wang et al., mRNA vaccine-elicited antibodies to SARS-CoV-2 and circulating variants. Nature 592, 616–622 (2021).

8. F. Schmidt et al., High genetic barrier to SARS-CoV-2 polyclonal neutralizing antibody escape. Nature 600, 512–516 (2021).

9. A. J. Greaney et al., Complete Mapping of Mutations to the SARS-CoV-2 Spike Receptor- Binding Domain that Escape Antibody Recognition. Cell Host Microbe 29, 44–57.e49 (2021).

10. L. Witte et al., Epistasis lowers the genetic barrier to SARS-CoV-2 neutralizing antibody escape. Nat Commun 14, 302 (2023).

11. G. Mantus et al., Evaluation of Cellular and Serological Responses to Acute SARS-CoV-2 Infection Demonstrates the Functional Importance of the Receptor-Binding Domain. J Immunol 206, 2605–2613 (2021).

12. J. E. Bowen, et al., SARS-CoV-2 spike conformation determines plasma neutralizing activity elicited by a wide panel of human vaccines. Sci Immunol 7, eadf1421 (2022).

13. D. F. Robbiani et al., Convergent Antibody Responses to SARS-CoV-2 Infection in Convalescent Individuals. bioRxiv 10.1101/2020.05.13.092619 (2020).

14. T. F. Rogers et al., Isolation of potent SARS-CoV-2 neutralizing antibodies and protection from disease in a small animal model. Science 369, 956–963 (2020).

15. L. Liu et al., Potent neutralizing antibodies against multiple epitopes on SARS-CoV-2 spike. Nature 584, 450–456 (2020).

16. S. J. Zost et al., Rapid isolation and profiling of a diverse panel of human monoclonal antibodies targeting the SARS-CoV-2 spike protein. Nat Med 26, 1422–1427 (2020).

17. E. C. Holmes, The Emergence and Evolution of SARS-CoV-2. Annu Rev Virol 11, 21–42 (2024).

18. T. N. Starr et al., Deep Mutational Scanning of SARS-CoV-2 Receptor Binding Domain Reveals Constraints on Folding and ACE2 Binding. Cell 182, 1295–1310.e1220 (2020).

19. K. Muthuraman et al., Human antibody targeting of coronavirus spike S2 subunit is associated with protection mediated by Fc effector functions. J Virol 99, e0152325 (2025).

20. J. S. Low et al., ACE2-binding exposes the SARS-CoV-2 fusion peptide to broadly neutralizing coronavirus antibodies. Science 377, 735–742 (2022).

21. C. Dacon et al., Broadly neutralizing antibodies target the coronavirus fusion peptide. Science 377, 728–735 (2022).

22. W. Li et al., Structural basis and mode of action for two broadly neutralizing antibodies against SARS-CoV-2 emerging variants of concern. Cell Rep 38, 110210 (2022).

23. X. Sun et al., Neutralization mechanism of a human antibody with pan-coronavirus reactivity including SARS-CoV-2. Nat Microbiol 7, 1063–1074 (2022).

24. P. Zhou et al., Broadly neutralizing anti-S2 antibodies protect against all three human betacoronaviruses that cause deadly disease. Immunity 56, 669–686.e667 (2023).

25. D. Pinto et al., Broad betacoronavirus neutralization by a stem helix-specific human antibody. Science 373, 1109–1116 (2021).

26. P. Zhou et al., A human antibody reveals a conserved site on beta-coronavirus spike proteins and confers protection against SARS-CoV-2 infection. Sci Transl Med 14, eabi9215 (2022).

27. F. Ruiz et al., Delineating the functional activity of antibodies with cross-reactivity to SARS-CoV-2, SARS-CoV-1 and related sarbecoviruses. PLoS Pathog 20, e1012650 (2024).

28. M. Lilly et al., Re-infection with SARS-CoV-2 is associated with increased antibody breadth and potency against diverse sarbecovirus strains. mBio 17, e0361225 (2026).

29. A. A. Cohen et al., Mosaic nanoparticles elicit cross-reactive immune responses to zoonotic coronaviruses in mice. Science 371, 735–741 (2021).

30. A. A. Cohen et al., Construction, characterization, and immunization of nanoparticles that display a diverse array of influenza HA trimers. PLoS One 16, e0247963 (2021).

31. A. A. Cohen et al., Mosaic RBD nanoparticles protect against challenge by diverse sarbecoviruses in animal models. Science 377, eabq0839 (2022).

32. G. B. Hutchinson et al., Nanoparticle display of prefusion coronavirus spike elicits S1- focused cross-reactive antibody response against diverse coronavirus subgenera. Nat Commun 14, 6195 (2023).

33. A. C. Walls et al., Elicitation of broadly protective sarbecovirus immunity by receptor- binding domain nanoparticle vaccines. Cell 184, 5432–5447.e5416 (2021).

34. G. G. Hendricks et al., Computationally designed mRNA-launched protein nanoparticle immunogens elicit protective antibody and T cell responses in mice. Sci Transl Med 17, eadu2085 (2025).

35. C. Fan et al., Neutralizing monoclonal antibodies elicited by mosaic RBD nanoparticles bind conserved sarbecovirus epitopes. Immunity 55, 2419–2435.e2410 (2022).

36. C. Fan et al., Cross-reactive sarbecovirus antibodies induced by mosaic RBD nanoparticles. Proc Natl Acad Sci U S A 122, e2501637122 (2025).

37. M. Wiatr et al., Memory B cell development in response to mRNA SARS-CoV-2 and nanoparticle immunization in mice. Proc Natl Acad Sci U S A 123, e2527869123 (2026).

38. T. Zang et al., Heteromultimeric sarbecovirus receptor binding domain immunogens primarily generate variant-specific neutralizing antibodies. Proc Natl Acad Sci U S A 120, e2317367120 (2023).

39. F. Muecksch et al., Affinity maturation of SARS-CoV-2 neutralizing antibodies confers potency, breadth, and resilience to viral escape mutations. Immunity 54, 1853– 1868.e1857 (2021).

40. F. Schmidt et al., Plasma Neutralization of the SARS-CoV-2 Omicron Variant. N Engl J Med 386, 599–601 (2022).

41. S. Cele et al., Omicron extensively but incompletely escapes Pfizer BNT162b2 neutralization. Nature 602, 654–656 (2022).

42. R. Pajon et al., SARS-CoV-2 Omicron Variant Neutralization after mRNA-1273 Booster Vaccination. N Engl J Med 386, 1088–1091 (2022).

43. C. Gaebler et al., Severe Acute Respiratory Syndrome Coronavirus 2 Neutralization After Messenger RNA Vaccination and Variant Breakthrough Infection. Open Forum Infect Dis 9, ofac227 (2022).

44. J. M. Carreño et al., Activity of convalescent and vaccine serum against SARS-CoV-2 Omicron. Nature 602, 682–688 (2022).

45. M. A. Tortorici et al., Repeated COVID-19 vaccine boosters elicit variant-specific memory B cells in humans. Cell Rep 45, 117052 (2026).

46. C. O. Barnes et al., SARS-CoV-2 neutralizing antibody structures inform therapeutic strategies. Nature 588, 682–687 (2020).

48. Z. Feng et al., A highly divergent cryptic SARS-CoV-2 lineage exhibits strong receptor binding and immune evasion. medRxiv 10.1101/2025.11.18.25340485 (2025).

48. M. A. Tortorici et al., Persistent immune imprinting occurs after vaccination with the COVID-19 XBB.1.5 mRNA booster in humans. Immunity 57, 904–911.e904 (2024).

49. F. Schmidt et al., Measuring SARS-CoV-2 neutralizing antibody activity using pseudotyped and chimeric viruses. J Exp Med 217 (2020).

50. A. Cho et al., Anti-SARS-CoV-2 receptor-binding domain antibody evolution after mRNA vaccination. Nature 600, 517–522 (2021).

51. Z. Wang et al., Memory B cell development elicited by mRNA booster vaccinations in the elderly. J Exp Med 220 (2023).

52. Z. Wang et al., Memory B cell responses to Omicron subvariants after SARS-CoV-2 mRNA breakthrough infection in humans. J Exp Med 219 (2022).

